# An ancient split of germline and somatic stem cell lineages in Hydra

**DOI:** 10.1101/2023.07.04.546637

**Authors:** Chiemi Nishimiya-Fujisawa, Hendrik Petersen, Tracy Chih-Ting Koubková-Yu, Chiyo Noda, Shuji Shigenobu, Josephine Bageritz, Toshitaka Fujisawa, Oleg Simakov, Satoru Kobayashi, Thomas W. Holstein

## Abstract

In many animals, germ cell segregation occurs during early embryogenesis to protect the genome, but its origin in basal metazoans is controversial. Here, we show in the freshwater polyp *Hydra* by clonal analysis and transgenic animals that interstitial stem cells comprise two separate stem cell populations, i.e., germline and multipotent somatic stem cells. We isolated genetically labelled stem cells for a global transcriptome study and discovered a broad set of germline-specific/enriched genes including *Prdm9, Pax5, Dmrt1*. In an alternative splicing analysis, we identified many genes with germline-specific isoforms; among them, male-specific isoforms of *Dmrt1* and *Snf5*. The somatic interstitial stem cell lineage was characterized by numerous neuronal control genes like *Neurog*. But all stem cells in *Hydra* also share a core of stemness genes that has its roots in unicellular eukaryotes. This suggests an evolutionary scenario in which, at the emergence of animal multicellularity, there was an early split into a stable germline and different somatic stem cell lineages.

## Introduction

Multicellularity has evolved independently in animals and plants. Both kingdoms also differ in the way they specify their germ cells. In plants, no germline exists and germ cells are specified lifelong by meristematic stem cells that also produce other somatic cells ^1,2^. But in many animals, germline cells, i.e., cells that are restricted to gamete production, are segregated from somatic cells at embryonic stages either by maternally inherited germline determinants or by induction of signals that suppress the somatic fate. The segregation of germline cells from the soma was the basis of the famous germline concept put forward by August Weismann at the end of the nineteenth century ^3,4^. Weismann studied the formation of germ cells (*Keimzellen*) in hydrozoans and he described stem cell-like interstitial cells as precursors to the germ cells (i.e., *Urkeimzellen* or germline cells) ^3^. His concept had a major impact on the modern synthesis of evolutionary biology, because it was limiting heredity to the germline, which forms a barrier to prevent accumulating mutations in somatic cells from entering the germline ^4,5^.

The Weisman “doctrine” was criticized and questioned in the past ^5–9^. At the heart of the argument against it were observations made in cnidarians, planarians, and sponges, indicating that similar to plants, germ cells arise continuously and life-long from stem cells competent to both somatic and germ cells ^5,7,8,10,11^. Evidence also emerged from expression analyses of genes that have a germline function in bilaterians (e.g., *vasa, nanos*, and *piwi*): those genes were expressed in stem cells of cnidarians and other basal metazoans that can give rise to somatic cells as well ^12–21^. These studies have led to the hypothesis of a “primordial stem cells” with an ancestral multipotency program thus questioning the idea of a germline in basal metazoans ^7^.

Here, we present long-term data testing the validity of the germline concept in the freshwater cnidarian *Hydra. Hydra* polyps reproduce also asexually (budding) and have been used to study longevity and regeneration ^22–25^. *Hydra* is also well suited to study stem cells (Fig. 1a) ^26–32^. There are ectodermal and endodermal epithelial stem cells, and interstitial stem cells ^28,33,34^. The prevailing view holds that interstitial stem cells differentiate somatic cells such as nerve cells, nematocytes, and gland cells, but also continuously segregate germline progenitor cells with a limited mitotic capability (Fig. 1a) ^26–28^. The sex of germline progenitor cells is autonomously determined and can be recognized by the growth pattern in asexual animals ^35,36^. However, when interstitial cells were eliminated ^37,38^ these germline cells did survive for several years, suggesting that they represent germline stem cells (GSCs) by its own ^27,35,36,39,40^. These findings have challenged the hypothesis that interstitial stem cells in *Hydra* are primordial stem cells ^7^ and suggests the existence of an independent germline stem cell (GSC) lineage.

**Figure 1.**
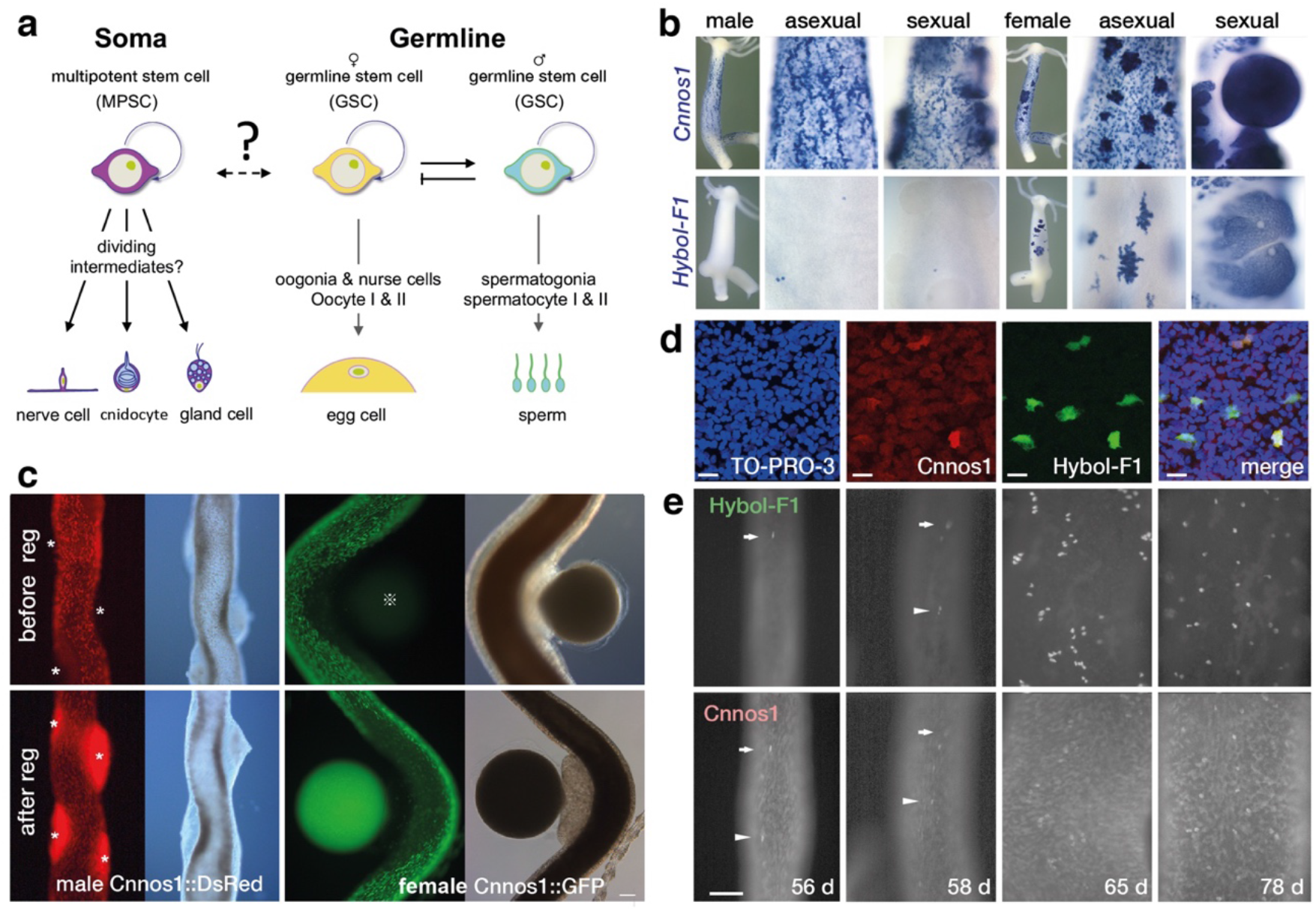
*Hydra* germline and somatic multipotent interstitial stem cells. (**a**) Somatic and germline stem cells in *Hydra*. (**b**) ISH expression analysis of *Cnnos1* and *HyBol-F1* in sexual polyps of *H. magnipapillata*, strain nem-1 (male and female) demonstrates that *HyBol-F1* is a female specific marker (occasionally a few cells express *HyBol-F1* in males but become eliminated by masculinization ^30^). (**c)** GSCs regeneration takes place from transgenic MPSCs only after removal of pre-existing GSCs (see text for details). (**d**) Double transgenic strain demonstrates HyBol-F1::GFP in GSCs and Cnnos1::DsRED expression in GSCs and MPSCs, counterstain with DNA marker To-Pro-3. (**e**) *in vivo* analysis of the genetic switch of double labelled (HyBol-F1::GFP / Cnnos1::DsRED) interstitial cells from MPSCs to GSCs. Animals were cut into small pieces to remove GSCs, cultured and analysed daily. The remaining MPSCs (DsRED^+^) started to produce single DsRED^+^/GFP^+^ female GSCs at 54-56 days after the onset of tissue regeneration (see experimental scheme in Supplementary Fig. 2). The clonal growth of two individual switched GSCs is indicated by an arrow (cell 1) and an arrow head (cell2) between 56d and 58d. Note that “switchable” cells display an increased expression of Cnnos1 compared to MPSCs with a rapid increase in number until 65d.

By using genetically labelled *Hydra* cell lines and a long-term tracing of interstitial cells, we can unequivocally demonstrate the existence of independent germline stem cells (GSCs). Contrary to previous assumptions, interstitial stem cells comprise two independent populations, (i) somatic interstitial multipotent stem cells (MPSCs) that develop only towards the neuronal and secretory pathways and never give rise to germline under a normal condition and (ii) germline stem cells (GSCs) ^27^. Despite the progress in defining *Hydra* cell type trajectories ^20,41^, the molecular identity and the properties of GSCs currently remain unclear. We therefore performed a detailed functional analysis of germline and somatic stem cells using specific genetic markers. Our FACS based study of *Hydra* stem cells provides an in-depth analysis of the transcriptome of a basal metazoan with intrinsic sex determination. In *Hydra* germ line specification, cell-type specific alternative transcription splice-sites play a critical role. Furthermore, we identified an archaic core cluster of epigenetic and transcription factors (TFs) in germ line and somatic stem cells that is shared throughout metazoan evolution by many stem cell systems up to human hematopoietic stem cells. Our analysis also sheds light on the evolution of animal germline and somatic stem cells and their implications for understanding the evolution of animal cell types.

## Results

### Germline stem cells constitute an independent cell lineage, MPSCs are somatic stem cells

In order to analyse the functional and molecular interrelationships of interstitial cell populations in *Hydra*, we genetically labelled interstitial cells and their differentiation products with DsRED or GFP by using the promoters of the cell type specific marker genes *Cnnos1, Dm5/Dmrt1, HyBolF1 and Hym176B* (Supplementary Table 1). *Cnnos1,* one of the two *Hydra nanos* genes, is a common marker for interstitial cells ^12,42^ (Fig. 1b, c). *HyBolF1* is a female-specific *Hydra* ortholog of an animal-specific family of RNA binding proteins (boule/DAZ/DAZL) that are exclusively expressed in germline cells and required for the meiotic progression of germ cells ^43,44^ (Fig. 1c, Supplementary Table 1, see below). Transgenic polyps were mosaic after hatching and must have been selected for polyps with more transgenic cells until polyps are homogenously labelled ^45^ (Supplementary Fig. 1, see also Materials and Methods). When we induced sex in homogenously labelled Cnnos1::GFP or Cnnos1::DsRED transgenic polyps, we surprisingly found that transgenic cells were completely absent from the sites of gamete formation (Fig. 1b, *upper row*) although endogenous *Cnnos1* was expressed in the sites (Fig. 1c) ^26,27^. We made this observation in four different strains (Supplementary Table 1), which were all cultured over years, but even repeated sexual induction cycles never resulted in transgenic gonads. This indicated that interstitial cells in *Hydra* form two distinct cell lineages, somatic multipotent stem cells (MPSCs), which do not give rise to germ cells, and germline stem cells (GSCs), which have unlimited mitotic capability by their own, and both lineages never mix.

We next used two strains (*Silv24* and *sGrn4,* for details see Supplementary Table 1) to test whether transgenic MPSCs can give rise to GSCs in GSC-depleted animals ^46,47^. GSC-depleted polyps were stochastically generated by cutting animals into small pieces that could be regenerated into polyps (Supplementary Fig. 1b-d). When regenerated polyps were sexually induced, 4 of 108 polyps of the male strain (*Silv24)* and 7 of 111 polyps of the female strain (*Grn4*) produced gonads that were strongly and uniformly labelled with no evidence of mosaicism (Fig. 1b *lower row;* Supplementary Fig. 1). This demonstrates that all GSCs in these polyps have been newly transformed from transgenic MPSCs in the absence of GSCs.

We analysed this surprising finding in more detail by using a double transgenic strain that additionally contained the exclusive germline marker *HyBolF1* (Fig. 1d). We crossed *HyBolF1::GFP* females with a F1 strain of the regenerated *Cnnos1::DsRED* males (Supplementary Fig. 2a; Supplementary Table 1). In the resulting female polyps, GFP was expressed in GSCs, and DsRED in GSCs and MPSCs (Fig. 1D, Supplementary Fig. 2a). When we cut out small pieces of tissue lacking GFP^+^ cells (Supplementary Fig. 2b), we found that DsRED labelled MPSCs started to express GFP quite some time after onset of tissue regeneration (Fig. 1e). Live cell imaging also demonstrated that then GFP^+^ cells then appeared at several sites of the gastric region and formed rapidly growing clones (Fig. 1E). These GFP^+^ cells were GSCs (*HybolF1*::*GFP*^+^; *Cnnos1*::*DsRED*^+^) that were transformed from MPSCs (*HybolF1*::GFP^−^/*Cnnos1*::*DsRED^+^*). This observation was remarkable since if MPSCs were competent to differentiate into GSCs, and repressive signalling from GSCs merely suppressed the transformation, the transformation should have occurred immediately after the removal of the GSCs. Clearly, this was not the case. No transformation occurred until the regenerated polys have grown to budding-ready sizes (ca. 2 mm long), which were much larger in size than newly hatched primary polyps (0.5 mm long) that already have GSCs (from the observation of *blF1::GFP* transgenic embryos). It sometimes took over 50 days after the tissue cut-out (Fig. 1e). Our observation indicates that MPSCs are not *per se* competent to differentiate into GSCs, but that transformation of MPSCs requires additional previously unknown signals. We anticipate that this transformation reflects the high regenerative and genome reprograming capacity of cnidarians.

The clear separation of the MPSCs and GSCs was unexpected and it raises the important question of how and when GSCs form during embryogenesis. We observed that the first GSCs (*BlF1::GFP*) only appeared in hatching primary polyps, while the first *Cnnos::GFP* cells appeared already during gastrulation. The identity of early *Cnnos::GFP* cells is unclear: they could be “primordial stem cells” from which MPSCs and GSCs form ^7,8,48^. Alternatively, GSCs do arise independently from MPSCs during embryogenesis. Histologic data on *Hydra* embryogenesis indicated that interstitial cells have two origins: ectoderm and endoderm ^49^. Possibly, GSCs occur in the endoderm and MPSCs in the ectoderm of *Hydra*. The observations done so far do not permit to differentiate between both options. However, our experiments uncovered that MPSCs are always somatic stem cells, and GSCs represent an independent and lifelong stable germline lineage that is already present in the hatching primary polyp (see Discussion).

### Sex reversal of GSCs

The existence of a germline in *Hydra* also implicates that the frequently observed sex-reversal of polyps occurs at level of GSCs but not of MPSCs as it was postulated in several articles Martin, 1997 #6039}^46^. Since MPSCs have been never observed to transform into GSCs under normal conditions, it is likely that female GSCs can switch into male GSCs and vice versa ^27^.

To test this directly, female or male GSCs were cloned (Supplementary Fig. 3). Strains with transgenic male or female GSCs could be easily generated by transplanting non-transgenic wild type AEP strains on top of polyps containing double-labelled female or male GSCs (Supplementary Fig.3a, b). Such chimeras contained transgenic GSCs without transgenic MPSCs (Supplementary Fig. 3c). To analyse sex reversal, small chimeric tissue pieces containing a variable number of transgenic MPSCs were cut out (Supplementary Fig. 4a). Only pieces containing a single transgenic GSC were selected (Supplementary Fig. 4b-g). They regenerated and were independently raised (clones of 10-20 polyps). After sex induction, all regenerates that were judged to have no transgenic cells produced testes without fluorescence (Supplementary Fig. 4h). In regenerates with transgenic cells, we analysed the frequency of sex reversal of male and female GSCs (Supplementary Fig. 4i, j). Of 20 independent animal clones, 19 clones produced both GFP^+^/DsRED^+^ eggs and GFP^-^/DsRED^+^ testes (Supplementary Fig. 4i). In clones with male GSCs (GFP^-^/DsRED) we did not observe any female GSCs (Supplementary Fig. 4j). These results decisively indicate that female to male reversal occurs at the level of GSCs. Our data are also consistent with observations in wildtype *Hydra vulgaris* AEP where female polyps can regularly switch into male polyps, but not vice versa.

### Transcriptome of Hydra stem cell lineages

The finding that GSCs and MPSCs constitute two independent, functionally different stem cell lineages, raises the question of their molecular identity. We therefore analysed the transcriptomic profiles of GSCs and MPSCs and compared them with neuronal cells and ectodermal and endodermal epithelial cells from the gastric region. Since asexually propagated polyps have the same genome, we have established several genetically independent clones of *Hydra* to average the biased gene expression that the use of only one clone could cause. Labelled cells were isolated by FACS (see Material and Methods, Supplementary Fig. 5, Supplementary Tables 1, 2). Hym176B neurons were the reference for fully differentiated cells of the MPSC lineage (Supplementary Fig. 6) ^50^. Cells of the GSC- and MPSC-lineage allowed us to monitor the down-regulation of the stem cell genes and the upregulation of cell type specific marker genes. The epithelial cells of the ectoderm and endoderm of the gastric region are two further independent stem cell lineages with differentiation potential and self-renewal capacity, and constitute the stem-cell niche for GSCs and MPSCs ^28,30,33^. Our RNA-seq analysis was performed by paired-end sequencing. A tSNE analysis of 110,303 high quality transcripts in our de novo transcriptome assembly revealed a clear separation of the different cell types (Supplementary Fig. 7). By mapping to the genome of *Hydra vulgaris* strain *magnipapillata* 105 ^24^ and *Hydra vulgaris* strain AEP ^41^, we obtained a coherent transcriptome profile between the two analyses. In addition, we identified 89 unique alternatively spliced genes including 6 event types when comparing GSC with MPSC via aligning to the AEP genome (Table 1, Supplementary Tables 11 and 12).

**Table 1.** Corresponding gene numbers and overlapping percentage of Spladder identified alternative splicing event types, adjusted p-value < 0.0001.

| Alternative splicing event type | MPSC vs ♀ GSC, N | MPSC vs ♂ GSC, N | ♀ GSC vs ♂ GSC, N | MPSC vs ♀ GSC<br>MPSC vs ♂ GSC<br>overlap, N | overlap / (MPSC vs ♀ GSC), % | overlap / (MPSC vs ♂ GSC), % |
| --- | --- | --- | --- | --- | --- | --- |
| 5' splice site | 8 | 10 | 1 | 3 | 37.5 | 30.0 |
| 3' splice site | 0 | 3 | 0 | 0 | NA | 0.0 |
| exon skip | 50 | 44 | 0 | 34 | 68.0 | 77.3 |
| intron retention | 10 | 1 | 0 | 1 | 10.0 | 100.0 |
| multiple exon skip | 17 | 10 | 0 | 7 | 41.2 | 70.0 |
| mutual exclusive exon | 0 | 2 | 0 | 0 | NA | 0.0 |

MPSC versus ♀ GSC: alternative spliced genes between MPSC and female GSC cell types.
MPSC versus ♂ GSC: alternative spliced genes between MPSC and male GSC cell types.
♀ GSC versus ♂ GSC: alternative spliced genes between female and male GSC cell types.
Overlap: number of overlapping genes between ♀ GSC and ♂ GSC comparisons.

In our global analysis, we identified 28,198 differentially expressed RNAs (Supplementary Table 2) (see Materials and Methods for detail). In a k-Means clustering (Fig. 2a) we found three clusters (labelled *AR/BG/BH*) that had very low expression in differentiated cells (nerve cells). Two of them (*BG*/*BH*) had their highest expression level in MPSCs and GSCs, one (*AR*) also in epithelial cells (Fig. 2a). Considering that gastric epithelial cells in *Hydra* are proliferating stem cells, we hypothesize that these clusters represent a repertoire of stem cell genes that are expressed in *Hydra’s* germline and somatic stem cells.

**Figure 2.**
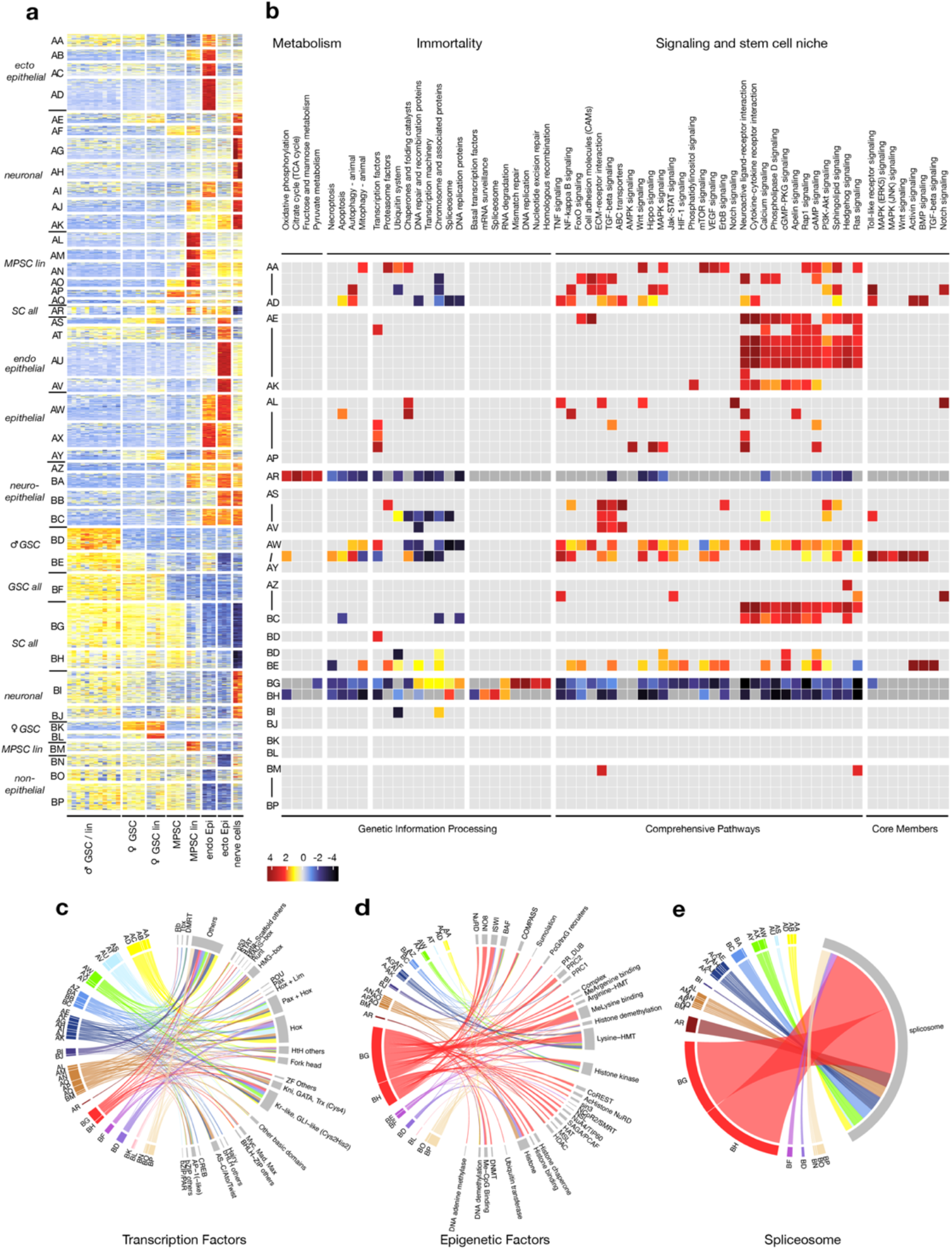
Cluster analysis of *Hydra* cell type specific genes. (**a**) *k-Means* cluster analysis of 21,283 differentially expressed transcripts from ten FACS isolated cell types revealed at a fold-change (FC) cut-off of 4 an enrichment of transcripts in 42 clusters (AA-BP). (**b**) KEGG based pathways analysis demonstrates a strong depletion in all pathways for the BG/BH and AR cluster except for the enrichment in DNA replication and repair, RNA metabolism and spliceosomes (note that database identifiers of KEGG pathways have been omitted clarity). **c**) Distribution of transcription across *Hydra* cell types by chord diagrams reveal an enrichment of *BG/BH* cluster (red) for Krueppel-/Gli-like zinc finger and Helix-loop-helix/bZIP-domain factors. (**d**) Epigenetic factors of the *BG/BH* cluster are enriched for Polycomb, ISM, BAF (**e**). Spliceosome factors are also enriched in the *BG/BH* cluster.

A KEGG enrichment analysis across all cell types and for all clusters (Fig. 2b) revealed that the *BG, BH,* and *AR* clusters were enriched with transcripts related to DNA repair/replication, RNA processing/synthesis, and metabolism, respectively. These three clusters were highly depleted in transcripts for apoptosis, autophagy, and cell differentiation (signalling pathways and cell type specific TFs) (Fig. 2b), whereas the BG/BH cluster was characterized by a particularly high content of factors acting in splicing and epigenesis (Fig. 2d, e). The two GSC clusters common to male and female (*BE/BF*) were enriched for transcripts related to DNA repair and recombination, chromosome and associated proteins, cell signalling including Wnt and TGF/BMP pathways, and proteasome and mitophagy factors. The MPSC cluster (*AP*) was enriched for transcription and cell signalling factors including those for neuroactive ligand-receptor interaction, AMPK, and Hippo signalling pathways. The lack of signalling factors in the common stem cell clusters (*AR*/*BG*/*BH*), and the enrichment of these in MPSCs (*AP*) and common GSC clusters (*BE*/*BF*) indicate that MPSCs and GSCs do not depend on the same signalling systems, but MPSCs and GSCs utilize unique signalling systems each other for the maintenance and differentiation (see below). Thus, MPSCs and GSCs are independent populations also at gene expression level.

### Genetic repertoire of germline stem cells

Our global cell type analyses revealed many genes that were expressed across several cell types (Fig. 2a). We therefore performed further two k-Means cluster analyses in which we compared the expression profiles from MPSCs versus male/female GSCs, and female versus male GSCs, regardless of whether they were also expressed in other cell types. We selected genes that showed a more than four-fold difference in both comparisons (Fig. 3, Supplementary Tables 3-6). A striking feature of this catalogue of germline factors was a remarkable similarity to that from PGCs in mammals ^51,52^. Here, we identified numerous orthologs of genes functioning in the mammalian germline determination cascade, from the epigenetic and transcriptional factors to signalling factors. Thus, our transcriptomic analysis was able to exhibit the evolutionary conserved module in the formation and maintenance of GSCs.

**Figure 3.**
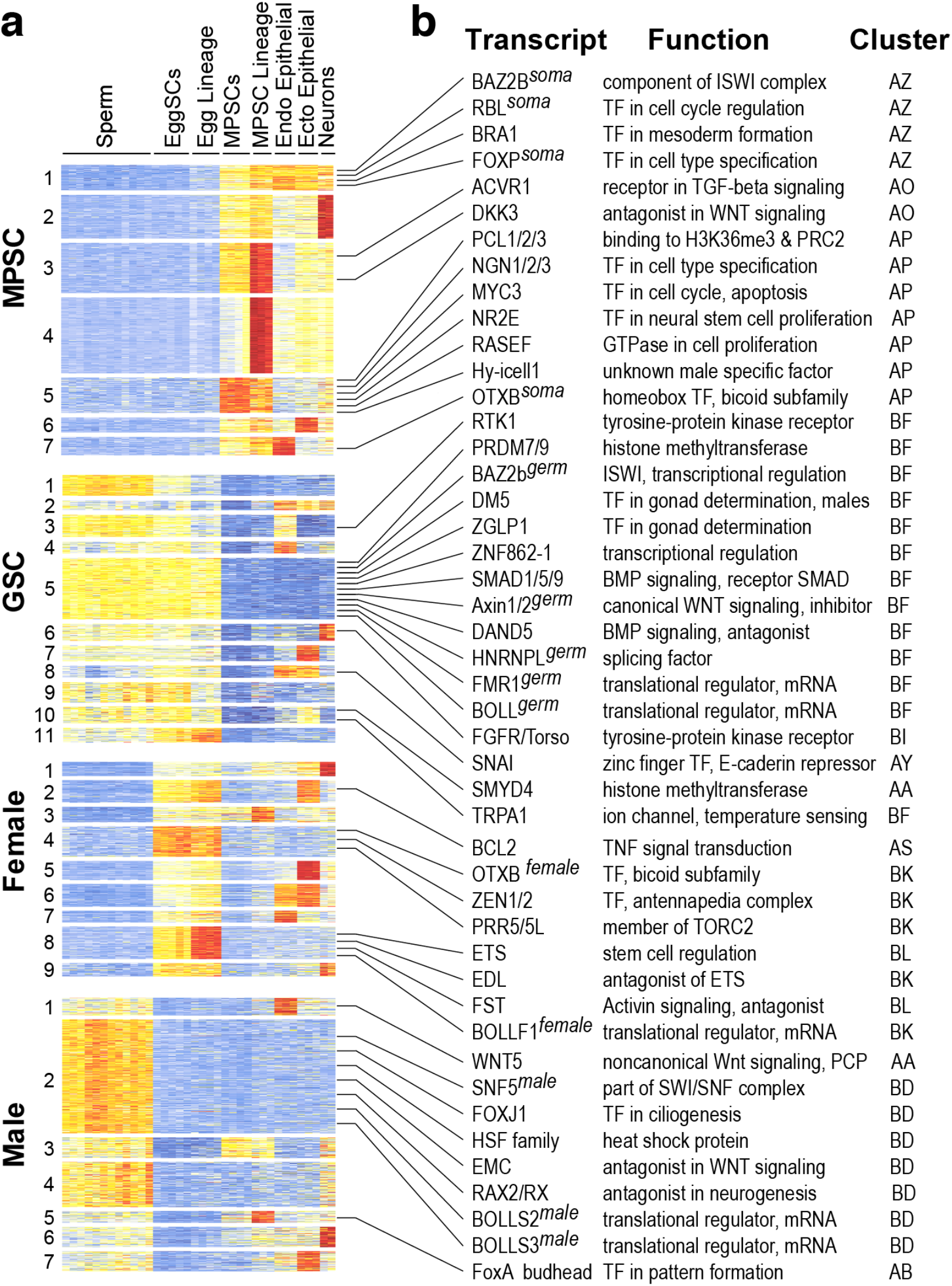
Cluster analysis of *Hydra* MPSCs, GSCs, and sexes. (**a**) *k-Means* cluster analysis of transcripts from FACS isolated MPSCs, GSCs, ♀ GSCs lineage and ♂ GSC lineage cells. (**b**) Selected transcripts and their distribution are depicted.

#### A putative germline function of Hydra Prdm genes

Prominent transcripts for epigenetic factors in the GSCs clusters encoded for PR domain zinc finger proteins (Prdms) ^51^ (Fig. 3). While *Prdm* genes are metazoan specific and have been identified from sponges to human ^53^. Their germline function was studied in detail in human and mice ^54–56^ ^57–59^. Here, Prdm1 (Blimp-1) and Prdm14 are the germline determinants that form a regulatory complex together with Pax5 and Oct4 to induce PGC formation and to repress somatic stem cell fates; *Prdm9* was expressed only in oocytes and spermatocytes, involving meiotic recombination. In *Hydra*, we found no direct orthologs of *Prdm1* and *Prdm14.* Instead, we found that *Prdm7/9-b* (DN4720) and *Prdm7/9-c* (DN1073) to be GSC specifically enriched. *Prdm7/9-a* (DN45181), *Prdm7/9-d* (DN1073), *Prdm-x1* (DN19633_c2_g1_i1), and *Prdm-x2* (DN37260) were upregulated in both MPSCs and GSCs and showed a similar expression pattern to *BG/BH* cluster genes. Considering that prdm14 has a stemness/pluripotency function, we expect these six Prdms expressed in GSCs and MPSCs have a similar function as mammalian Prdm1/9/14. Also factors upstream or downstream of Prdms in mammals are activated in *Hydra* GSCs. We therefore suggest that the entire Prdm-mediated network for specifying mammalian PGCs is also present in *Hydra*, from the Wnt/Bmp-dependent induction to Pax5-Oct4-mediated transcription ^52,60,61^. Other *Prdms* exhibited also a highly cell type specific expression pattern in *Hydra* (Fig. 3, Supplementary Table 6) further suggesting that *Prdm* genes are main cell type specifiers in *Hydra.* represent an evolutionary conserved core module that already specified germline cells in cnidarians.

#### TFs, RBPs, and epigenetic factors in Hydra’s germline

As additional important TF from mammals that we identified in *Hydra’s* germline were Zglp (*DN1230*) a TF with GATA-like zinc finger that determines oogenetic fate ^62^, Sohlh (*DN17539*) a HLH TF for spermatogenesis, oogenesis and folliculogenesis, and the mesodermal Zinc finger TF Snail (*DN33799i*) that is transiently upregulated during specification of PGCLC ^63^.

In addition to TFs, we found eleven *Boule/Daz/Dazl* genes encoding germline-specific RNA-binding proteins ^43,44^. Three of these are expressed in both sexes (*DN28822; DN65338; DN17508)*, three are enriched in the male germline (*DN49425; DN11241; DN29260*), and five in the female (*DN20680; DN57674; DN69704; DN70567; DN20303*). It is unclear why *Hydra* has so many *Boule* genes, but the expression of these genes seems to be dependent on the developmental stage of the germline cells. The expression of *Boule* genes shared by both sexes was restricted to GSCs, and those genes may be involved in the identity/maintenance of GSCs under cnidarian conditions where gonads do not exist.

Despite of numerous similarities between *Hydra* and mammalians described above, we did not detect expression of *Tfap2* (also known as Ap2-gamma) in GSCs of adult polyps, which was described as the critical regulator of germ cell induction in mammals and the related hydrozoan *Hydractinia* ^48,64^. However, we identified *Tfap2* expression in early embryos during gastrulation and in adult polyps in endodermal epithelial cells (*DN35822*) (*DN35822*). This suggests that this gene is required in *Hydra* for embryonic germline-specification or/and for endoderm specification.

Unlike in mammals, we found no obvious evidence of an involvement of FGF signalling in GSC control ^65^. Since *Hydra* GSCs are located in an “open” niche, where stem cells move around, we expected GSC density homeostasis to occur by FGF signalling, such as in the mammalian testis, but found no evidence of this. Fifteen transcripts encoding FGF ligands were all enriched in epithelial cells, some of which are also in GSC lineage and none in the MPSC lineage. The four FGFRs (*DN30102; DN26686; DN42200; DN47446*) were expressed mainly in epithelial cells as well, showing no significant difference between GSCs and MPSCs. Thus, FGF signalling is seemingly not involved in *Hydra* GSC density homeostasis so far. Whether some of the RTK genes up-regulated in GSCs (*DN45201_c2_g2_i2; DN22690; DN42060_c0_g1_i5; DN46687_c2_g1_i3/4/10; DN30084*) have such a function, remains to be shown.

We also found the following GSC genes in *Hydra*: *Trp* (Transient Receptor Potential Cation Channel) genes (DN44558; DN46148_c4_g4_i1/2/4; DN47478; DN46858; DN47877) ^66,67^, some of which can sense the environmental temperature as Thermo-Trp, and may trigger temperature-dependent sexual differentiation in *Hydra* ^68,69^; *Smyd4c* (DN43878), one of the three *Smyd4* genes in *Hydra*; *GRM* (Glutamate Metabotropic Receptor) genes (DN37977; DN46238). Two genes reported as female germline specific (*Hyfem-1; Hyfem-2*) ^20^ were also expressed in male GSCs in our hands.

### Sex-determining factors in GSCs

Sexual dimorphism is common to all multicellular organisms suggesting that sex determining mechanisms of germline cells can be traced back to ancestral unicellular eukaryotes. However, it remains unknown to what extent these mechanisms are conserved in animal evolution. One reason for this is that is, in bilaterians, the sexes in the germline are instructively determined by the gonads, which evolved independently among species after the evolution of mesodermal tissue. In cnidarians, which have no mesoderm, sex determination of GSCs occurs in a cell-autonomous manner ^21,26^, and they might have retained the ancestral mechanism. It is therefore a key question, how the sex of GSCs can be determined in *Hydra*. Our sex-specific germline transcriptomes enabled us to identify the sexual genes to a depth that was previously not possible for any prebilaterian animal (Figs. 2 and 3; Supplementary Tables 3-5). When we compared male and female GSCs, we found further striking examples for alternative splicing, sex specific genes, and many male specific transcription factors (Fig. 3; Supplementary Tables 3-5).

To unambiguously analyse the splicing patterns between the cell-specific transcripts of GSCs and MPSCs, we also mapped the RNA-seq reads to *Hydra vulgaris* strain AEP ^41^ and systematically tested for cell type specific splicing variants. Using this approach, we detected >100 conserved cell type-specific alternative splice variants between germline and somatic stem cells. Table 1 shows that male and female specific germline stem cells have a similar splicing profile to each other, but differ significantly from MPSCs. This pattern was found for most splice types (Supplementary Figure 9), although only alternative 5’prime sites (TSS) were found between male and female GSCs. More than half of the splicing events overlap between male and female GSCs when compared with MPSCs (Table 1).

Among the TFs genes enriched in female GSCs we identified *Otx2 ^female^ (DN38831_c0_g1_i1)* as a major factor. Otx2 is a mammalian key regulator for neuronal differentiation, but also for the segregation of the germline from the soma in response to Bmp4 ^94^. The splice variant *Otx2^female^* was strongly upregulated only in female germline cells, while *Otx2^soma^*(*DN38831_c0_g1_i3*) was upregulated only in somatic cells (MPSC lineage and in endodermal epithelial cells) (Supplementary Figure 10 and Supplementary Tables 2, 6 indicating the female specific function of *Otx2 ^female^*. We also found *zerknuellt (Zen) (DN28821_c0_g1_i1*) also known to act downstream of Bmp2/4 ^95^; a function in the context of germline cells was unknown so far. Two isoforms of *Edl* (*ets-domain lacking*) (*DN37558_c0_g2_i1, DN37558_c0_g2_i2*) were also expressed in the female germline, suggesting an antagonizing role against the *Ets* (Yamada et al., 2003) during oocyte/nurse cell development (Supplementary Table 4).

One example for sex specific isoforms was Snf5 (also called smarcB1), a core component of the SWI/SNF/BAF chromatin remodelling complex. This complex is also a critical regulator for metazoan cell type differentiation, and a mutation of *Snf5* causes cancers in mammals ^70,71 72^. The canonical isoform *Snf5 ^univ^* (*DN34020_c0_g1_i1*) ^73 74^ was enriched in female GSCs and MPSCs, while the non-canonical isoform *snf5^male^* (*DN34020_c0_g1_i2*) was restricted to male GSCs and SpLCs (Supplementary Tables 3-5 and Supplementary Figure 10b, female GSC shared the same splicing pattern as MPSC (data not shown). Interestingly, the two differentially expressed N-terminal exons are similar, suggesting that the non-canonical isoform arose by exon duplication Supplementary Figure 10. Moreover, the genome configuration of the *Snf5* gene is highly conserved among cnidarians examined so far (*Clytia, Hydractinia*, and *Nematostella*). We were able to further confirm the male-specific expression of the exon on the genome browser of *Clytia*, where transcriptomes from sexual male and female medusae were mapped on the genome ^75^. Based on the conserved genome organization and expression among Cnidaria, we presume that the *Snf5^male^* has a specific function in males rather than it is a dominant negative form. Thus, we propose that the male-specific *Snf5* variant is involved in the sex determination of germline cells also in other cnidarians, either upstream or downstream of Dm domain genes.

As distinct male-determining gene we identified the *Hydra* ortholog to bilaterian DM domain genes from insects *Doublesex (Dsx)* and *Dmrt1* from vertebrates Dsx/Dmrt1 is an evolutionary conserved transcription factor that is expressed in testes and other sexually dimorphic cells, and acts in primary bilaterian sex determination. In *Drosophila* it is expressed in the somatic gonadal primordium and subject to alternative splicing in male and female flies; in vertebrates it is expressed in the somatic cells of the embryonic gonad, the Sertoli cells of the testes, and male premeiotic germ cells ^76,77^. The elevated expression of *Hydra Dm5*/*Dmrt1* (*DN20040_c1_g2_i1*) in male GSCs and their lineage, the half dose expression in female GSCs and its decline in the female GSC lineage (Supplementary Table 4) suggests its ancestral sex-determining function in *Hydra* male GSCs. That *Dm5*/*Dmrt1* was enriched at a lower level in female GSCs also suggests a potential function in *Hydra*’s sex-reversal of GSCs ^26,47^. As reported in Anthozoa, one gene (*DN16498*) was expressed in differentiating female germline cells (nurse cells) ^78^. More *Dm/Dmrt/Dsx* genes were expressed somatically in neuronal cells and/or in epithelial cells. Importantly, our discovery that GSCs themselves do express *Dmrt1* in *Hydra* instead of a somatic gonad tissue – as in bilaterians – indicates that sex determination by *Dmrt1-like* genes has evolved in cnidarians in unicellular eukaryotes (see Discussion).

Among the epigenetic factors we found a large number of alternatively spliced GSC-specific isoforms. A major factor is *Hnrnpl^germ^*(DN46875_c0_g1_i1), which itself is involved in splicing, an integral component of ISWI complex that has a short AT-rich extra exon that can be involved in its translational control, and *Ldb* variants (LIM domain-binding protein; DN29902_c0_g1_i4; _c0_g1_i2), both of which lack the conserved N-terminal (Supplementary Figs. 9, 10); *Baz2b^germ^*(*DN41468*_c0_g1_i1) and Supplementary Figure 10b, female GSC shared the same splicing pattern as MPSC (data not shown) (Supplementary Figs. 9, 10 and Supplementary Table S12).

### Interstitial MPSCs are restricted to neuronal and secretory cells

The direct comparison between GSCs and MPSCs revealed many MPSC enriched genes that were also identified in our global analysis (*AO, AP,* and *AZ*) (Figs. 2b and 3). Among the TFs we identified important pluripotency-related transcription factors known from insects and vertebrates ^79^ (e.g., Brachyury, Fox-P, Otx2, and Rbl). We also identified MPSC enriched signalling molecules (e.g., *Activin-Receptor1, Dkk3*, and *Rasef*) and epigenetic factors, e.g., of the ISWI complex and *Polycomblike* (*Pcl*) (Figs. 2B and 3) (Supplementary Tables 7, 10). Many differentiation factors in this lineage were completely absent in the GSCs-lineage (Fig. 3) or had GSC- and MPSC-specific splice variants (e.g., *Baz2b, Neurogenin, Otx2, Pcl, Rbl*) (see above, Supplementary Fig. 9, 10). When we directly compared the expression levels of MPSCs and MPSC-lineage cells we could also identify genes that exhibited higher expression levels in MPSCs than in the MPSC-lineage cells. Out of a total of 56 TFs that were expressed in both cell types, we found three MPSC-enriched TFs from the AO/AP clusters, i.e., *Neurogenin* (*DN14758_c0_g2_i1*) (a master TF that determines the fate of neurogenesis), *Nr2e* (nuclear receptor) (*DN35411_c1_g1_i1*), and three paralogs of the Yamanaka factor Myc, i.e., *Myc1* (*DN35396_c0_g1_i2*), *Myc2* (*DN12617_c0_g1_i1),* and *Myc3/MycL* (*DN65524_c1_g4_i1*). Other major Yamanaka factors were orthologs of *Oct4/Pou5* (*DN34464*); *Sox2* (*DN39389*) (*BF* cluster), and *Kruppel/Gli* (*DN65575*). Co-enriched in GSCs (e.g., in the *BG* or *BF* cluster) were *Myc2, Sox2, Oct4/Pou, and Gli/Kruppel*. *Hydra Myc1* and *Myc2* have been previously described to be specifically activated in the rapidly proliferating cell types of the MPSCs lineage and in proliferating gland cells ^80,81^. All other TFs, such as *Atonal, Ascl*, and *SoxB* exhibited a higher expression level in MPSC-lineage cells or even in nerve cells. Also, the transcription factor *FoxO* (*DN40663_c0_g1_i2*), which has been described as the factor for stem cell maintenance and longevity in *Hydra* ^82,83^ had its strongest enrichment in MPSC/MPSC lineage cells and neurons. The high number of neuronal TFs enriched in MPSCs and MPSC lineage cells raises the question of the identity of this somatic stem cell lineage. If MPSCs were primordial stem cells, one would expect a much broader representation of stemness related TFs. It was also obvious that most members of the “conserved germline multipotency program” like piwi and vasa were much stronger expressed in GSCs than in MPSCs. We therefore assume that MPSCs do not have the properties of primordial stem cells, but represent a stem cell population in *Hydra* restricted to neuronal/neurosecretory cells.

We tested this hypothesis by further increasing the resolution of our already sensitive bulk RNA-seq analysis by a scRNA-seq analysis on isolated DsRED-labelled MPSCs and MPSC-lineage cells (strain *Silv24reg*, Supplementary Table 1). By this approach we could visualize continuous changes in the differentiation pathways of the MPSC transcriptome with highest precision. Figure 4A shows the cluster analysis of scRNA-seq data from isolated DsRED-labelled MPSCs and MPSC-lineage cells and for comparison their distribution in bulk RNA. A remarkable outcome of this scRNA-seq analysis of isolated MPSCs was those genes of the AR and *BG/BH* stemness cluster had their highest expression level in the least differentiated MPSCs (cluster 5, orange) and strongly decreased along the MPSC differentiation paths. Here, we identified one path towards neuronal cells and one towards nematocytes and gland cells. We did not find a path towards transcripts specific for GSCs or GSC lineage cells including *Tfap2* transcripts ^48^ (Fig. 4b). The expression of genes of the MPSC lineage clusters indicates, however, a neuronal (*AE-AK*) and nematoblast/nematocyte (*AL-AP* / *AZ-BC*) differentiation path. Both developmental paths are in accord with the recently found trajectories in a scRNA-seq analysis of whole animals ^20,41^. These data therefore support our main findings which have led to the conclusion that MPSCs do not give raise GSCs in intact animals and that MPSCs represent a somatic stem cell population largely committed towards neuronal neurosecretory differentiation. Genes of the *BG/BH* cluster, however seem to represent the core cluster for stemness genes

**Figure 4.**
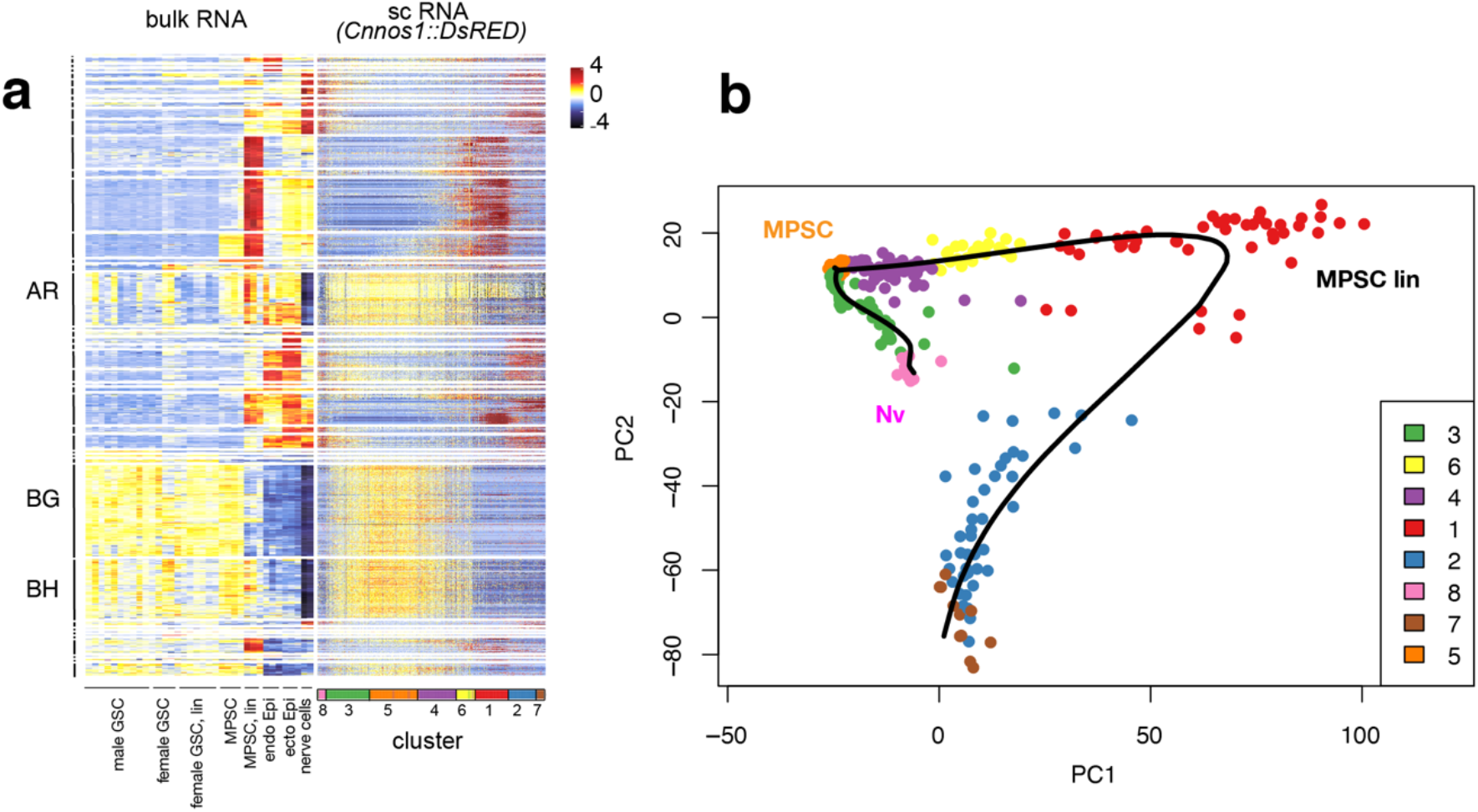
Single cell sequencing of MPSCs. (**a**) Cluster analysis of transcripts from bulk RNA and scRNA isolated DsRED-labelled MPSCs and MPSC-lineage cells (strain *Silv24reg*). Note the distribution of *AR, BG* and *BH* clusters and the clustering of sc-transcripts. (**b**) t-SNE representation of MPSCs indicates continuous changes in the differentiation pathways with two major branching developmental paths in MPSC differentiation, the neuronal (3, 8) and the nematoblast/gland cell directed line (1, 2, 4, 6, 7). Note that genes of the *AR* and *BG/BH* cluster had their highest expression level in the least differentiated MPSCs (5) and decreased along the two differentiation paths.

### Ancestry of stem cells and early origin of the germline

The *BG/BH* stemness cluster contained 2212 transcripts and revealed genes controlling basic stem cell functions as replication, DNA repair, transcription, translation and RNA processing (Figs. 2; Supplementary Tables 7-10). On the level of TFs, we found stem cell related TFs (i.e., Gli- and Kr-like and Zf factors) and factors that have a function in regulating the homeostasis between mammalian embryonic stem cells (ESCs) and germline cells ^79^ (Fig. 2c, Supplementary Table 10). Especially the *BG* cluster was rich in TFs involved in the maintenance of a pluripotency network in ESCs e.g. *Myc2, Lhx8, SoxB, Rbl1, PaxB1, p53, p63*, and *Tex10 (DN53200_c0_g1_i1), a* gene encoding for a Sox2-interacting protein that together with Tet1 and p300 sustains stemness in somatic and germ cells ^84,85^. The number of epigenetic and RNA-dependent factors (e.g., Polycomb, Trithorax group II, and the BAF/PBAF/ncBAF chromatin remodelling complex) was much higher in the *BG/BH* cluster than in any other cluster ^87,88,104–107^. It should be finally emphasized that our scRNA-seq and trajectory inference analysis revealed that genes of the *BG/BH* cluster were almost exclusively expressed in stem cells at the very beginning of the trajectories, but not in differentiating cells (Fig. 4). Thus, genes enriched in the *BG/BH* cluster seem to be of ultimate importance for maintaining the pluripotency of somatic and germline stem cells in *Hydra*.

In order to further unravel the linkage between germline and somatic stem cells during animal cell type evolution, we determined the transcriptome age index (TAI) ^86,87^ of *Hydra*’s GSCs/MPSCs, the corresponding cell lineages and of epithelial cells. Our phylostratigraphy analysis revealed three distinct age groups (Fig. 5a). The group with the oldest TAI included MPSCs, GSCs and GSC-lineage cells, the middle group were epithelial cells, and those with the youngest TAI nerve cells and MPSC lineage cells. This distribution corresponded to the contribution of the different phylostrata (*ps*) (Fig. 5b). Most MPSCs and GSCs specific genes evolved already at the level of unicellular eukaryotes (Fig. 5c) while sex-specific genes appeared later on the level of sponges (Fig. 5c). Genes of the MPSC-lineage including neurons as well as germ layer specific epithelial genes appeared only on the level of the Eumetazoa/Cnidaria (Fig. 5c). These data have important implications: The basic germ layers (i.e., ectoderm and endoderm) were an eumetazoan invention and the large set of neuronal genes appeared on the level of basal eumetazoans. With respect to the evolutionary origin of germline and somatic stem cells our data show that genes giving rise to sperm and egg cells were an invention of the basal metazoans, while the entire repertoire of stemness genes in GSCs and MPSCs is much older and can be traced back to basal eukaryotes. We also analysed how many of the *Hydra* stem cell genes were evolutionary conserved up to humans. We used genes of the *BG/BH* cluster and performed a cross-species analysis of orthologous *Hydra* genes against the transcriptomes of sponges ^16,18^, sea anemones ^18^, planarians ^88,89^ and mice (i.e., embryos and the adult hematopoietic system). We started with 2212 genes of the *BG/BH* cluster and discovered a strong coverage across the different phyla. 1345 *BG/BH* genes were also enriched in sponge archaeocytes, sea anemone precursor cells, and planarian neoblasts, and 931 of them in embryonic mice (prenatal stage 1-11) and early stages of adult mouse haematopoiesis (Fig. 5d). The strongest enrichment in mammals was found in the early myeloid, lymphoid and thymocyte hierarchy including bone marrow progenitor cells. These metazoan stemness genes revealed genes acting in cell cycle regulation, RNA metabolism, DNA repair and chromatin modification (Fig. 5e) and they also included an epigenetic stem cell core network with DNA-acetylation and -methylation, members of the SWI/SNF related complex and the complete component of PCR2 network (Fig. 5f). Thus, the major repertoire of epigenetic programming of *Hydra* somatic and germline stem cells is conserved throughout metazoan evolution, up to humans.

**Figure 5.**
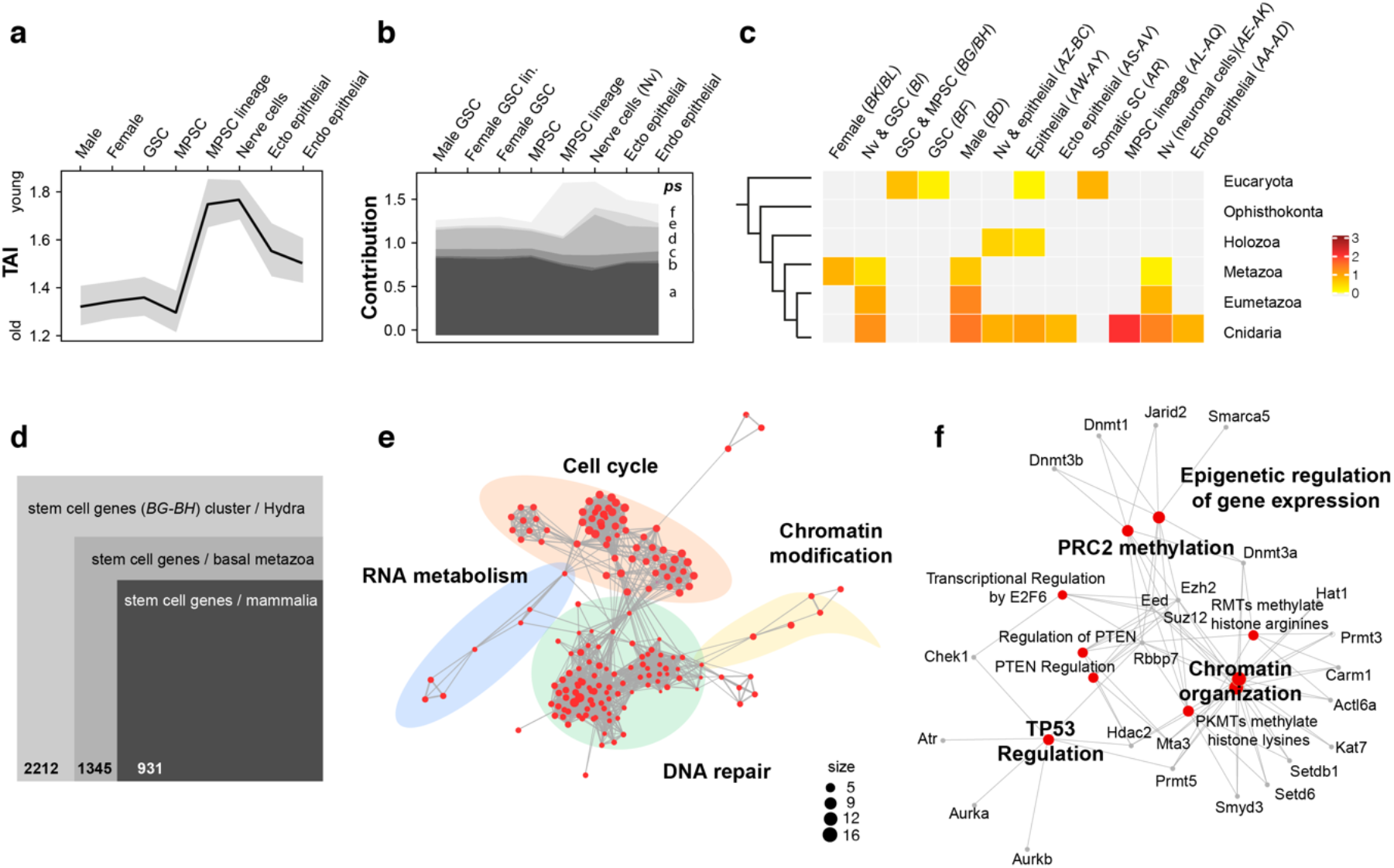
Evolutionary conservation of stemness factors from *Hydra* stem cells in metazoan evolution. **(a-c)** Transcriptome age profile of *Hydra* stem cells and phylostrata comparisons. (**a**) Cumulative transcriptome age index (TAI) for *Hydra* cell types reveals anciency of *Hydra* stemness genes. (**b**) Transcriptome indices split according to the origin of the genes from the different phylostrata, based on the same cell types as in (a). (**c**) Depiction of the analyzed phylostrata and enrichment of selected cell-type related gene clusters. The log2 of the enrichment factor are shown. (**d**) Diagram showing the overlap of the 2212 transcripts from the *BG/BH* stem cell cluster of *Hydra* with transcripts from basal metazoans (planarian, sponge, sea anemone) and mouse. 1345 genes are conserved in at least one basal animal and in mouse. (**e**) Enrichment map of reactome pathway enrichment analysis demonstrate a strong impact of DNA repair, RNA metabolism, cell cycle and chromatin modification in the function of conserved stem cell genes. (**f**) Gene-concept network of conserved stem cell genes involved in chromatin modification identifies members of the PRC2 with a conserved role in stem cell regulation.

## Discussion

In this study we have analysed germ cell formation in the freshwater polyp *Hydra* by using transgenic strains expressing germline specific markers. We could demonstrate by clonal analyses that in *Hydra* an independent lineage of germline stem cells (GSCs) exists that is separated from multipotent somatic stem cells (MPSCs) (Fig. 6). This is different from the current view questioning the existence of a germline in most basal metazoans ^7,90^. There, it was postulated that GSCs arise in basal animals by the transformation of somatic cells as in plants and all other multicellular eukaryotes (Fig. 6). In most animals, however, GSCs are formed by a germline that is sequestered during embryogenesis, either by epigenetic mechanisms or by maternal preformation (Fig. 6).

**Figure 6.**
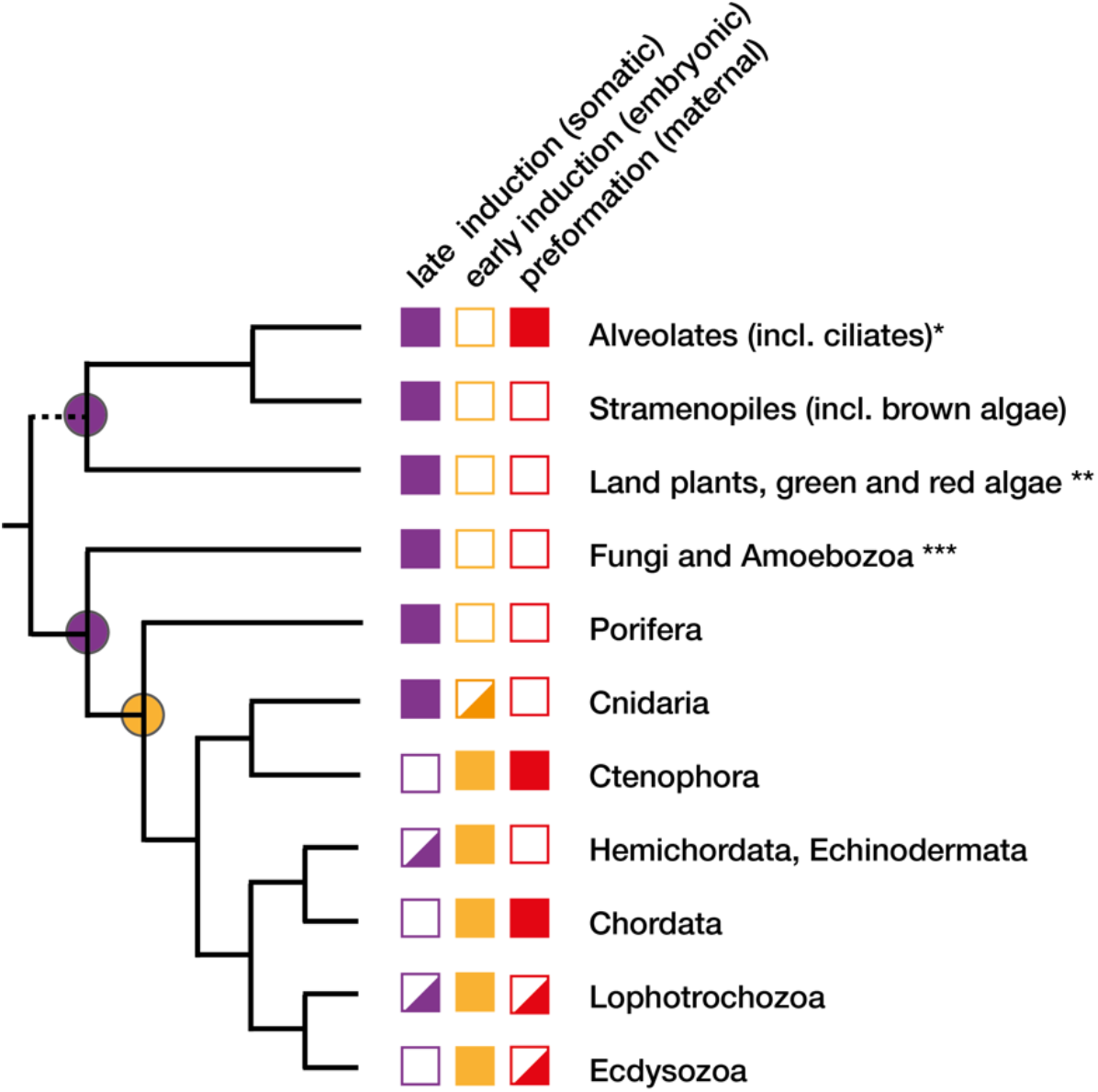
Mechanisms of germ cell specification across multicellular eukaryotes. Cladogram shows a simplified grouping of multicellular eukaryotes according to recent phylogenomic studies ^135, 136^. The distribution of germ cell specification mechanisms follows Whittle and Extavour (2017)^13^ and Buss (1983) ^1^and findings in this study. According to the life cycle of multicellular eukaryotes three modes of germ cell specification can be distinguished: a late induction from somatic cells (purple), an early induction during embryogenesis (yellow), and a maternal preformation (red). The presence of a trait in the sub-taxa of a kinship group is expressed by filling in the symbols, i.e. filled, split and open symbols indicate full or partial presence or absence. While only the preformistic sequestering of GSCs was considered in Weisman’s germ plasm theory ^6, 137^, an early separation of GSCs from somatic cells during embryogenesis by epigenetic mechanisms has a similar effect. In both cases germline cells are protected from mutations (variation) in somatic cells that cannot be transmitted to the germ line. This does not apply to protists, plants and fungi, which might have evolved different mechanism to protect the mutation rate of the germline, e.g., by (*) a silenced germline nucleus in ciliates ^138^. Mechanisms that are under discussion in plants (**) and fungi (***) are haploid selection ^139, 140^ and selective DNA strand segregation ^141^. It is proposed that the embryonic separation of a germline by epigenesis evolved only in the animal kingdom (filled yellow symbols), while mechanisms for somatic induction evolved repeatedly in multicellular eukaryotes (filled purple symbols) and in colonial metazoans (split purple symbols).

We isolated pure GSCs, analysed their transcriptome, and compared it with MPSCs and the differentiated cells. By this we uncovered for the first time a large set of germline- and sex-specific genes including *Pax5, Snf5*, and *Dmrt1* and genes of Wnt and Bmp signalling pathways as conserved constituent for a germline. In MPSCs we found many neuronal transcription factors e.g., *Neurog*, the master neural gene, suggesting that MPSCs in *Hydra* are neuronal stem cells. We also identified a cluster of epigenetic and transcription factors, which was enriched in germline and somatic stem cells and conserved throughout metazoan evolution. We also provide evidence that this archaic core of stemness genes originated in unicellular eukaryotes, remained stable during later evolution and become restricted to the stem cell lineages in multicellular animals.

An unexpected outcome of our transcriptome analysis was the broad overlap of *Hydra* GSCs genes with mammalian PGCs ^51,60,91^. Many factors of the cascade of germline determinants, from the epigenetic and transcriptional factors to the upstream acting signalling factors are highly conserved in *Hydra* GSCs suggesting that they represent an evolutionary conserved module in the formation and maintenance of GSCs. The fact that *Hydra* shares such a wide range of germline genes with vertebrates including mammals indicates that evolution of animal GSCs has a very deep origin in the evolution of the Metazoa. We anticipate that it can be traced back to the age of unicellular cellular animals. So far, no study has been reported yet ^92^, but we expect that GSCs had their origin in the sexual phase of unicellular animals which might express a similar set of germline genes like Hydra GSCs. We hypothesize that in unicellular animals, the germ cell and somatic cell are separated temporally: a single cell proliferates mitotically as a somatic cell at one time, while in other occasions, the same cell goes into miotic phase as a germ cell to produce gametes (Fig. 6). Thus, a similar set of common germline genes found in *Hydra* would be expressed during the sexual phase.

We also identified many genes that have a function in bilaterian animals in sex determination of the somatic cells in the tissue of the gonads. One example is the doublesex (*DM5/dmrt1)* gene, which has a main function in sex determination of insects and vertebrates, and which is mainly expressed in somatic cells of the gonad. This gene is strongly expressed in *Hydra* GSCs. We hypothesize that sex determination by the *dmrt1* gene initially originated from ancestral unicellular organisms that express a *dmrt1* gene both in the sexual and asexual phase in male cells (Supplementary Tables 11, 12). This sex-specific expression has been maintained up to *Hydra* where it performs a function in GSCs during sex-determination in a cell-autonomous manner. During bilaterian evolution, the expression must have been gradually restricted to somatic (mesodermal) gonadal cells, where it has a function in instructive sex determination. We also resume that this shift of pathways from GSCs in diploblastic animals to the mesoderm was tightly linked to the origin of triploblastic bilaterian animals ^93,94^.

The evolution of germ cell specification has recently been discussed in several revealing articles, but these mainly concern bilaterians and not pre-bilaterians ^5,9,95–97^. Our work was focused on pre-bilaterians. We show that *Hydra’s* GSCs display all features of a bilaterian germline and they share an archaic core set stemness genes with somatic stem cells. Given that Metazoa evolved from single-celled eucaryotes, in which germ and somatic potentials are expressed separately in time, the real innovation in the evolution of the germline was not the acquisition of a gametogenic lineage but rather the loss of gametogenic potential ^98^ and its substitution by somatic potential in somatic cells (Fig. 6). This is manifested by a multitude of bilaterian stem cell systems leading to an enormous variety of animal cell types. Thus, this ingenious division of labour has made somatic stem cells the determining factor for the fitness of genetic innovations that have emerged by mutations primarily in germline cells. By comparison, plants have no germ line and evolved with haploid selection and a high transposon tolerance a different mechanism to overcome the deleterious effects of somatic mutations (Fig.6).

We believe that the segregation of a germline from the soma is a fundamental property of Metazoa, but the hypothesis is hard to prove as many basal Metazoa have colonial growth patterns (e.g., in sponges or marine hydrozoans and corals) where GSCs can form in the adults (Fig. 6). This was shown recently in the colonial hydrozoan *Hydractinia symbiolongicarpus* where *Tfap2* acts in the commitment of germ cell fate of adult polyps ^48^. This is different to *Hydra* where *Tfap2* is only expressed during early embryogenesis. *Hydra* separated from colonial marine hydrozoans 215 Mya ago ^99^ and lives as a solitary polyp, although its asexual reproduction by budding is reminiscent of its former colonial way of life. Similar to *Hydractinia, Hydra* has also lost its Medusa generation, which is the sexual generation in most hydrozoans and scyphozoans with a complete life cycle, and which has been considered to be responsible for a significant gene loss in *Hydra* ^75,100^. It should be stressed however, that hydrozoans are extremely pleiomorphic with repeated loss of the polyp or medusa generation ^101,102^. In colonial anthozoans, a lower mutation rate for GSCs than for the somatic tissue was reported for coral *Orbicella faveolate* ^103^, which indicates the existence of a germ line in anthozoans. Thus, both scenarios are currently plausible for cnidarians: a repeated separation of the germline in different clades or a loss of the germline in colonial forms.

Is the selective benefit of the germline measurable? In a study on vertebrates that differed in their mode of germ cell determination (preformation by germ plasm *versus* induction), it was reported that the acquisition of germ plasma accelerates the rate of protein evolution ^96^. Although this hypothesis was refuted in another study with a larger pan-bilaterian dataset ^104^, the idea that the germline can unlock the potential for somatic innovation ^9^ is still attractive. It might therefore be worthwhile to include prebilaterian animals and plants in this type of comparison. We expect that there was a selective pressure on the formation of a germline even in the earliest metazoans (Fig. 6). This was also a starting point for the radiation of somatic stem cell systems in the last common ancestor of bilaterian animals ^105,106^.

A main damage of the genome during aging is the breakdown of heterochromatic domains and an increased expression of retrotransposons. Argonaut family proteins Piwi and Piwi-related and their associated Piwi-interacting RNAs (piRNAs) are critical factors to ensure heterochromatin maintenance and to downregulate retrotransposons ^107^. We found that the strong expression of *Hydra* Piwi/Piwi-related orthologs ^108,109^ (*BG* cluster) was correlated to a weaker expression of transposable elements (TE) and less family counts in the corresponding stem cell gene clusters (*AR/BG/BH*). Conversely, in clusters enriched for transcripts in nerve cell (*AE-AK*) and MPSC lineage cells (*AP, AQ*) we found high family counts for TEs. It will be interesting to clarify whether TE expression has a direct impact on genome integrity, e.g., whether it is linked to the genomic structure of stemness genes by using the promoter of nearby genes. Such a mechanism could enforce the genome integrity by Piwi-interacting microRNA system.

Our data also indicate how the cell type identity between GSCs and MPSCs might have evolved. GSCs and MPSCs are located in a shared stem cell niche, where they are exposed to the same extracellular signals. In this identical cellular environment, they have to establish their cellular identity in a cell-autonomous manner. They can do it by alternative splicing of the critical cell fate determining factors. However, at present the conservation of alternative splicing and/or alternative start sites of orthologous genes in cnidarians is unknown. Lessons from vertebrates suggest, however, that phylogenetic conservation is low. Only 10% of the conserved exons between mouse and humans display alternative splicing in both species, although a core set of ancestral mammalian splicing regulators was identified ^110–112^.

## Materials and Methods

### Animals

The *Hydra vulgaris* strain *magnipapillata* 105 ^113^ and *Hydra vulgaris* strain AEP ^49^ were used and cultured as described^113^. To induce gametogenesis, the AEP animals were starved for one week and then fed twice per week.

### Cloning and Constructs

For the various EGFP and DsRED reporter constructs, promoter fragments of Cnnos1, HyBolF1, Hym176B, and actin were amplified by PCR from genomic DNA of AEP animals and subcloned into the hoT G ^45^, which resulted in hoTG-Cnnos1-DsRED, hoTG-Cnnos1-eGFP, hoTG-HyBolF1-eGFP, HotG-Hym176B-eGFP, HotG-DM5, and HotG-Actin-eGFP. Derivative constructs were generated by restriction digest and subsequent subcloning into *Hydra* transgenesis vectors, pBSSA-AR or its derivative containing the *Hydra* hsp70 minimal promoter (pBSSA-AR-hsp.mini-EGFP), which contain the *Hydra* actin promoter and the RFP gene..

### Generation of Transgenic Hydra Polyps

Generation of transgenic *Hydra* polyps was carried out as previously described ^45^. To ensure whether transgenic cells have a transgene, we constructed and used a vector system for *Hydra* transgenesis described before ^132^. Each transgene construct was injected into fertilized eggs of the AEP strain. Hatched polyps were collected about 6 weeks after injection and maintained individually. Polyps ubiquitously expressing the transgene were generated by clonal propagation, asexual budding. For each transgene, several independent transgenic lines were obtained, and at least two lines were analyzed. Details of generation of transgenic lines are described in Supplementary Table 1.

### Cell type isolation

GFP^+^ or DsRED^+^ MPSCs were collected from the founder transgenic lines that have no labelled GSCs (see above, *DB115, -116, sGRN4* and *Silv24*) (*SI Appendix*, Fig. S1, Supplementary Table 1). Female GSCs were collected from either F2 strains (LB3, LB30, Ly25, see Fig. 1D) or chimeric strains containing solely GFP/DsRED double-positive GSCs (*LB3/AEP_D, Ly25/AEP_DG*) (*SI Appendix*, Fig. S4, Supplementary Table 1). Male GSCs were collected from chimeric strains containing solely GSCs expressing DsRED^+^ (*LB2/AEP_D, LB3/AEP_D, LB25/AEP_D, Gold93B*) (*SI Appendix*, Fig. S5, Supplementary Table 1).

### RNA-sequencing and transcriptomic data

For the bulk RNA-sequencing analysis the starting RNA material was 280 ng/sample. For the library preparation we used Illumina TruSeq stranded mRNA Kit and the Illumina HiSeq 1500 platform with 15-45 Million reads/sample. The RNA reads were first *de novo* assembled using Trinity ^114^ and aligned to the genome of *Hydra vulgaris* strain *magnipapillata* 105 ^24^ for most of the analyses in the paper. Later, the RNA reads were aligned to the newly published *Hydra vulgaris* strain AEP genome ^41^ using HISAT2 ^116^ for alternative splicing analysis. For the single cell transcriptomic analysis, the Drop-seq system was used following available protocols ^115^. Pre-processing of the single-cell transcriptome data was done following the Macosko computational cookbook. The reads were aligned to the genome of *Hydra vulgaris* strain *magnipapillata* 105 ^24^.

### Differential expression analysis

Differential expression analysis was performed with DESeq2, comparing all isolated and sequenced cell types against each other. Reports of fold change and significance values (padj) of this all-vs-all comparison were used to isolate significantly regulated transcripts.

### Semi-supervised k-mean clustering

For clustering and visualization of expression patterns, raw counts were normalized and transformed with DESeq2, using the rlog function. rlog values of significant regulated transcripts (padj = 0.01) with various minimal fold change cut-offs (256, 128, 64, 32, 16, 8 and 4) were extracted using an in-house written python script and further processed in R. After centring and scaling the transcript expression values an initial k-mean clustering was performed on transcripts with the highest minimal fold change of 256 (log2 fold change of 8). Because of these highly discriminative transcripts 36 high quality cell type specific clusters could be manually determined and used as “core clustering”. With decreasing fold change cut-offs, the additional transcripts were sequentially added to these “core cluster” using semi-supervised clustering, implemented in the Vegclust package. For each round of semi-supervised clustering one extra cluster was added, to consider for cell type specific expression pattern only visible for transcripts with lower fold change between samples. These steps finally resulted in 42 distinct clusters, for transcripts with a minimal fold change of 4 between at least one cell type comparison (DESeq2 analysis^117^, TPM analysis^118^, Supplementary Tables 2-10).

### Ortholog identification and Kegg enrichment

*Hydra* orthologs from mammals (human, mouse) were identified with *OrthoFinder* using proteome resources using default settings. Based on these relationships KEGG annotations were transferred to *Hydra* sequences using an in-house written python script. Here, available KEGG annotations from the whole ortholog group were transferred. KEGG enrichment analysis was performed with *ClusterProfiler in R* using default conditions. The code was slightly modified not only to detect enriched but also depleted KEGG terms. All clusters from each step (fold change threshold) of the semi-supervised k-mean clustering were tested for enrichment and depletion. As a summary of this analysis relevant enriched/depleted KEGG terms were manually selected and the log2 fold change was visualized in a heatmap using *ComplexHeatmap in R*. Here, always the highest value was chosen within the enrichment results between the different fold change steps. The development of the enrichment / depletion within the various fold change steps are shown in the supplements visualized in a heatmap.

### Chord Diagram

The chord diagram was produced in R using the “cyclize” package. Transcripts were either automatically annotated with KEGG (transcription factor) or manually using BLAST (epigenetic factors). Shown are only deferentially regulated transcripts with a minimal fold change of 4.

### Transcriptome age index

Transcriptome age index analysis were performed in R using the *myTAI* package. Transcripts were annotated with phylostrata based on our ortholog analysis. The contribution diagram was visualized with ggblot2 and the enrichment analysis were done with *ClusterProfiler*.

### Cross species comparison

To compare expression of *Hydra* stem cell genes in other organisms, quantitative cell type specific expression data of sponge, *Nematostella*, planarian and mouse were used. The raw expression matrices of scRNA-seq data from sponge, sea anemone and planarian were processed in R to consistently normalize the data using the *Scran* and *Scater* packages. For bulk RNA-seq from sponge and mouse, already normalized data were used. All data were centered and scaled separately. Only *Hydra* stem cell genes which have orthologs in at least one basal animal (sponge, *Nematostella* and planarian) and in mouse are considered to be conserved and used in this analysis. Therefore, orthologs from *Hydra* stem cell genes in sponge, *Nematostella*, planarian and mouse were identified from our *OrthoFinder* analysis using pairwise orthologs. In a first selection process, the expression pattern of these conserved genes was compared in the basal animals. For this comparison the expression matrices were merged on the basis of the *Hydra* ortholog and clustered with k-means, to identify only genes which are highest expressed in the basal animal stem cells. These selected genes were then translated in ENSEMBLE mouse ids and used to extract mouse developmental and anatomy expression data from *Genevestigator*. From these main categories, quantitative data from mouse embryonic development, embryonic cell types and hematopoietic stem cell system were used for comparison. Here, the datasets were merged based on the gene id and again clustered with k-means to identify genes with highest expression in mouse stem cells.

### Reactome enrichment

In order to use the ENSEMBL mouse ids for Reactome pathway analysis, the ids were translated in ENTREZ ids using the bitr function in the *ClusterProfiler* package in R. Reactome pathway analysis was done with the Reactome package using default settings. The enrichment results were visualized in an enrichment Map which organizes enriched terms into a network with edges connecting overlapping gene sets. The category size was set to ##. Genes involved in the enriched pathways were visualized with the gene-concept network function. Here, only pathways and the corresponding genes around the PRC2 members Ezh2, Suz12, Eed, Rbbp7, Dnmt3a were selected.

### Alternative Splicing Analysis

The SplAdder package ^119^ was used to systematically analyse alternative splicing events between different cell types, including MPSC versus female GSC, MPSC versus male GSC and female GSC versus male GSC. Six event types were identified by the program, including alternative 5’ splice site (alt_5prime), alternative 3’ splice site (alt_3prime), exon skip (exon_skip), intron retention (intron_retention), multiple exon skip (mult_exon_skip) and mutual exclusive exons (mutex_exons). We used adjusted p-value ≤ 0.0001 as cut-off and the boundary cases were manually curated by checking the read profiles in the *Integrative Genomics Viewer* (IGV).

## Data and materials availability

Data are available in the main text or the supplementary information The original raw data and a comprehensive Supplementary Table 2 of all annotated transcripts have been submitted to GEO (GSE163910).

## Acknowledgments

We thank Suat Özbek for critically reading various versions of the manuscript. Funding: This work was supported by grants to C.N-F. (KAKENHI #15K14565 from JSPS), to S.K. (KAKENHI #25114002 from MEXT; KAKENHI #24247011 from JSPS), to T.W.H. (DFG-SFB 873-A1-2/3, DFG-SFB 1324-A5-1/2, and CHS foundation) and to a joint proposal (DFG/FWF D.A.CH) to O.S. and T.W.H.

## Author contributions

C.N.-F. designed the work, established transgenic strains (A14, A16, DB115, DB146, sGrn4, Silv.24, Silv24regenerate, Silv24regenerate -6, LWA, LWA-4, LWA-16, LB2, LB3, LB28, LB30, Gold93B, LY25, LB3 / AEP_DG, LY25 / AEP_DG, LB2 / AEP_D, LB3 / AEP_D, LY25 / AEP_D), cultured strains, performed transplantation and stem cell cloning assays, isolated RNA and prepared cDNA libraries. FACs analysis was performed by C.N., S.S., and H.P. H.P. designed, set up and performed bioinformatic analyses except the de novo assembly of the AEP transcriptome and differential expression analysis. J.B. performed scRNA sequencing of MPSCs. C.N.-F., T.C.-T.K.-Y., T.W.H., and O.S. performed alternative splicing analyses. O.S. and T.W.H. performed transposon analysis. T.F. analyzed neuronal data. S.K. designed and supervised the initial work. C.N-F., H.P., O.S. and T.W.H. analyzed the data and wrote the manuscript.

