## Supplementary Information for "An ancient split of germline and somatic stem cell lineages in Hydra"

**This PDF file includes:**

Figures S1 to S10
Table S1
Legends to Table S2 to S12

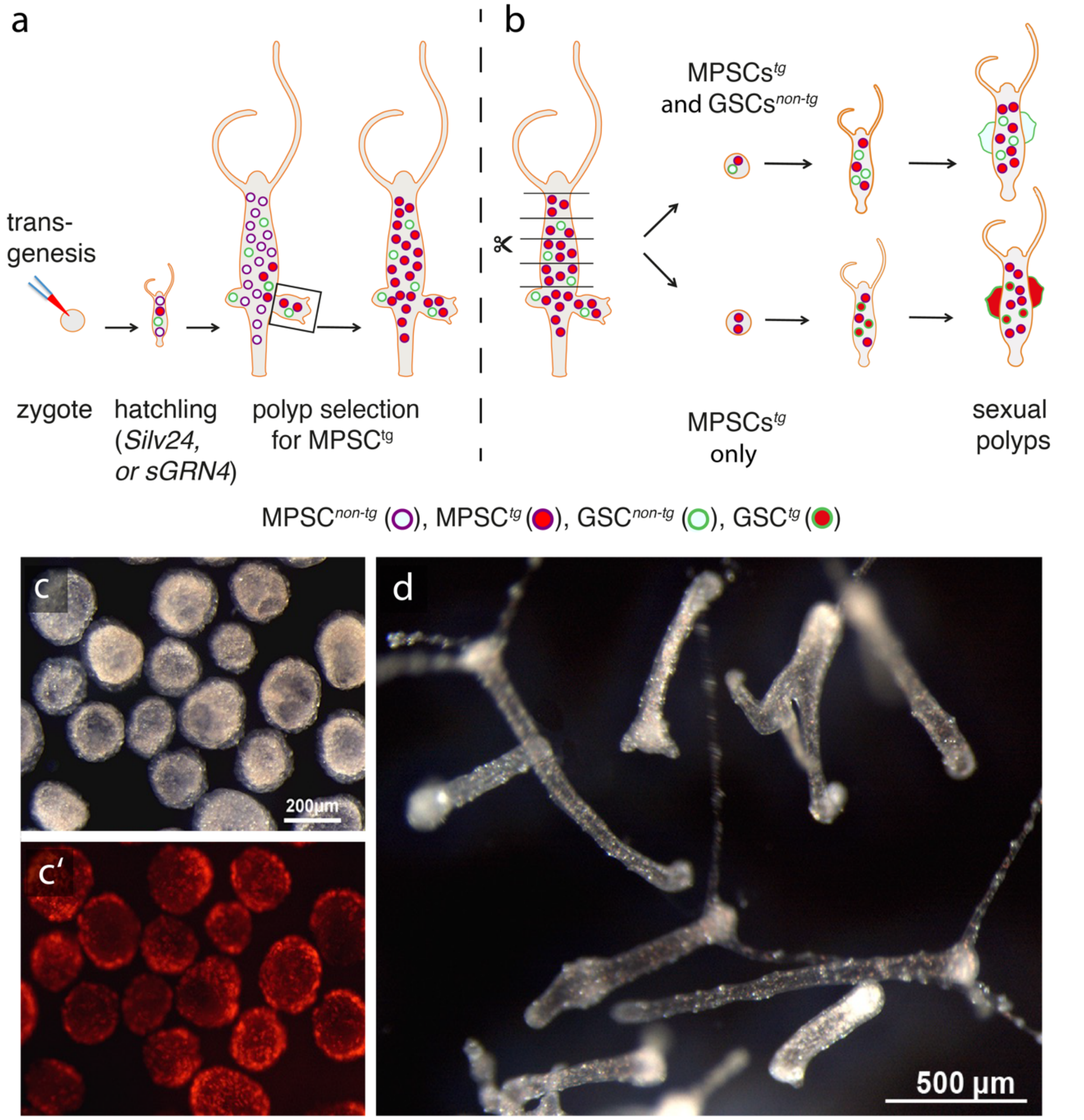


**Fig. S1.** Experimental scheme to generate strains with transgenic MPSCs and transgenic GSCs. (a) Transgenic founder polyps listed in Supplementary Table 1 were produced by injecting marker constructs. Strain sGRN4 was labelled with *Cnnos1::eGFP* and strain Silv24 with *Cnnos1::DsRED*. After hatching, primary polyps (hatchlings) were mosaic for transgenic and non-transgenic MPSCs. Growing cultures were selected for polyps with more transgenic cells until all polyps were uniformly labeled. After sex induction, transgenic cells were absent from the sites of gonad formation indicating that GSCs are not labelled (Fig. 1b, see text). (b) Cutting the animals into small pieces generated pieces containing transgenic MPSCs (see also Supplementary Fig. 7 for isolating small pieces). After regeneration, polyps were induced to form gonads. Most gonads were not labeled, but some were, indicating that they were formed from transgenic GSCs, which in turn must have originated from transgenic MPSCs (see also the transformation assay in Supplementary Fig. 2). (c) Isolated small pieces exhibiting homogenous labelling of MPSCs with *Cnnos1::DsRED* (strain Silv24) (c’) and (d) regenerated small polyps that were grown into larger steady-state polyps.

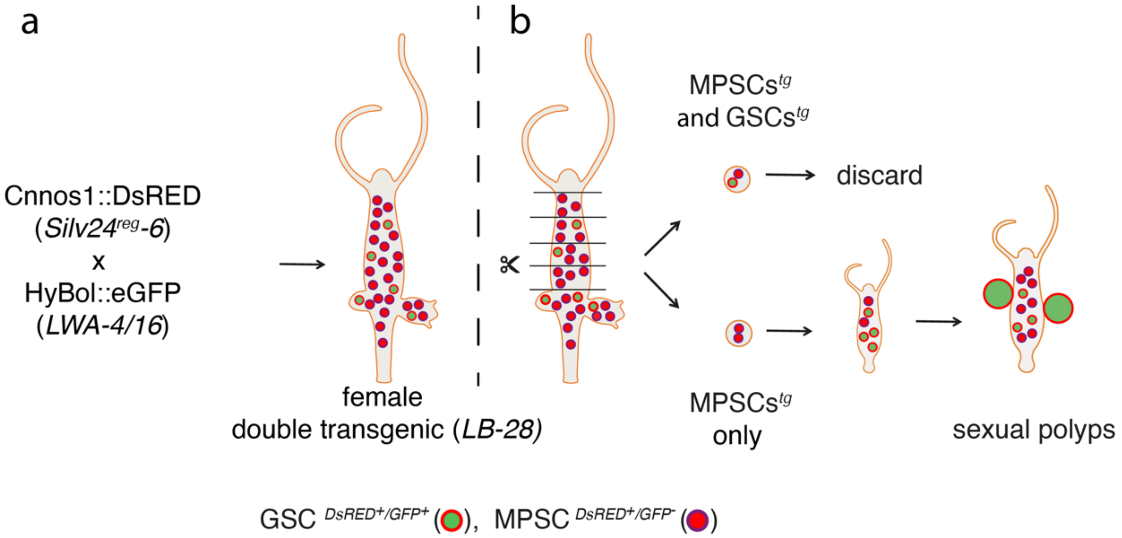


**Fig. S2.** Transformation assay for MPSCs using double-labelled transgenic animals. (a) To obtain transgenic animals that contained double-labelled GSCs, females of the strain LWA-4/16 (*HyBolF1::eGFP*) were crossed with males of the strain Silv24$^{reg}$ -6 (F1 strain of regenerated *Cnnos1::DsRED* polyps). HyBolF1 is a female GSC specific marker (see text). In the resulting strain (LB-28), female GSCs expressed eGFP under control of the *HyBol* promotor (*HyBolF1::eGFP*) and DsRED under control of the *Cnnos1* promotor (*Cnnos1::DsRED*). By comparison, MPSCs expressed only *Cnnos1::DsRED* (see also Fig. 1d, Supplementary Table S1). (b) By cutting the animals into small pieces we produced regenerates that contained DsRed labelled MPSCs and eGFP/DsRed doubled-labelled GSCs. Only pieces with DsRed labelled MPSCs were allowed to regenerate polyps. Regenerates were then screened for the appearance of double-labeled cells, i.e. cells expressing eGFP/DsRed. These cells must have originated from MPSCs that started to express the germline marker (*HyBolF1::eGFP)* (see Fig. 1e and text for details).

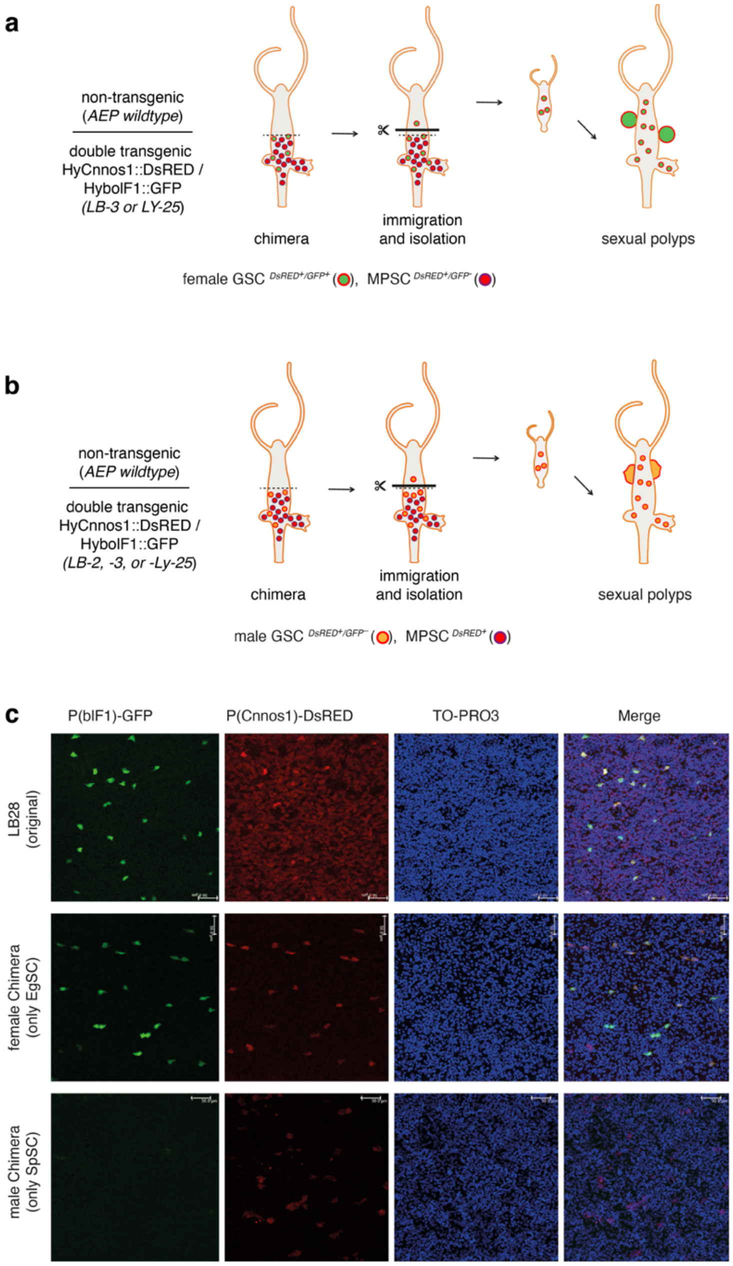


**Fig. S3.** Cloning of transgenic male and female GSCs by graft-migration method. Wild type AEP polyps were grafted on transgenic donor polyps and labelled GSCs migrated into the upper unlabelled AEP tissue, which was isolated again and regenerated intact chimeric polyps used for the isolation of GSCs by FACS (Supplementary Table 1). (a) For female GSC cloning, donor polyps contained double-labelled female GSCs (*Cnnos1::DsRED* and *HybolF1:eGFP)* and DsRED labelled MPSCs (*Cnnos1::DsRED*) (strain LB-3 or LY-25). (b) For male GSC cloning, donor polyps contained male GSCs (*Cnnos1::DsRED*) that did arise from primary polyps (strain Gold93B; not shown) or from switched female GSCs (strains LB2, LB3 and LY25). (c) Confocal images show the distribution of transgenic GSCs in the original transgenic female strain (LB-2, -3, -28, -30) and in the female and male chimera strains. The original strains contained numerous DsRED$^+$ MPSCs (*Cnnos1::DsRed*), while both chimera strains are free of any transgenic MPSCs. Male strains contain DsRED$^+$ GSCs that are eGFP Note that the GFP$^+$ female GSCs (*blF1::GFP*) are much stronger labelled with DsRED$^+$ than MPSCs.

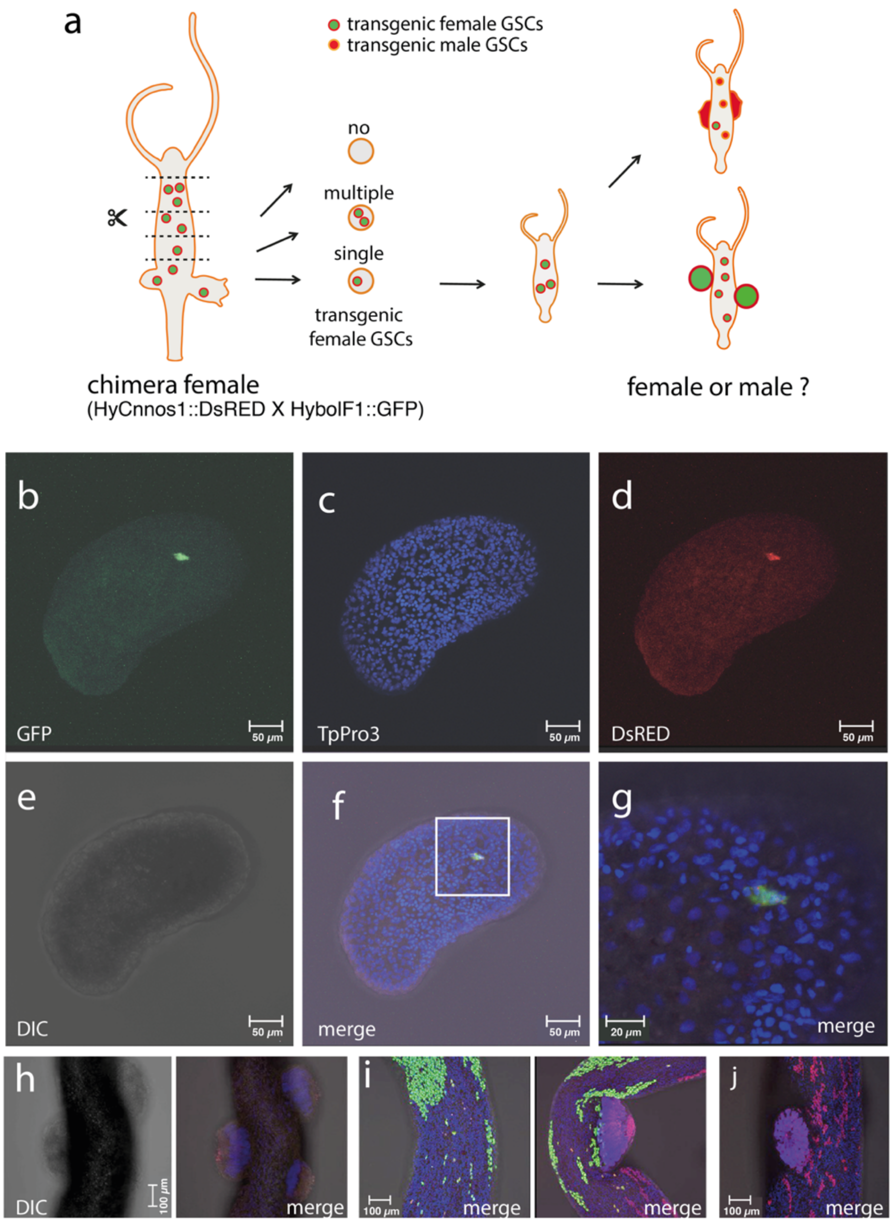


**Fig. S4.** Cloning of single GSCs. The potential of GSCs to reverse their sex was analysed as outlined (a). Female chimeric polyps (see Supplementary Fig. 4) with labelled female GSCs (*HyBolF1::GFP / HyCnnos1::DsRED double)* were cut into small pieces and raised to sexually mature polyps that were tested if they produced eggs or testes. Note, only pieces that after microscopic inspection, exhibited a single double-labelled female GSC were raised, and before sexual induction each polyp had to form a clone of 10-20 polyps. (b-g). A small piece containing a single double labelled female GSC expressing eGFP (*HyBolF1::eGFP*) (b) and DsRED (*Cnnos1::DsRED*) (d). Small pieces were counterstained with nuclear stain (TO-PRO3) (c, f, g) and additionally inspected with DIC (e). (g) close-up of the region with the single double-labelled cell shown in (f). (h-j) Sex reversal in animals arising from a single GSC. Tissues classified as GSC free developed testes without fluorescent cells (h) In 19 of 20 clones from pieces with one female GSC, GFP+ egg spots and RFP+ testes form simultaneously, $GFP^{+}$ egg patches and RFP+ testes form concomitantly (i). In clones from pieces with male GSCs, no female patches were observed (j).

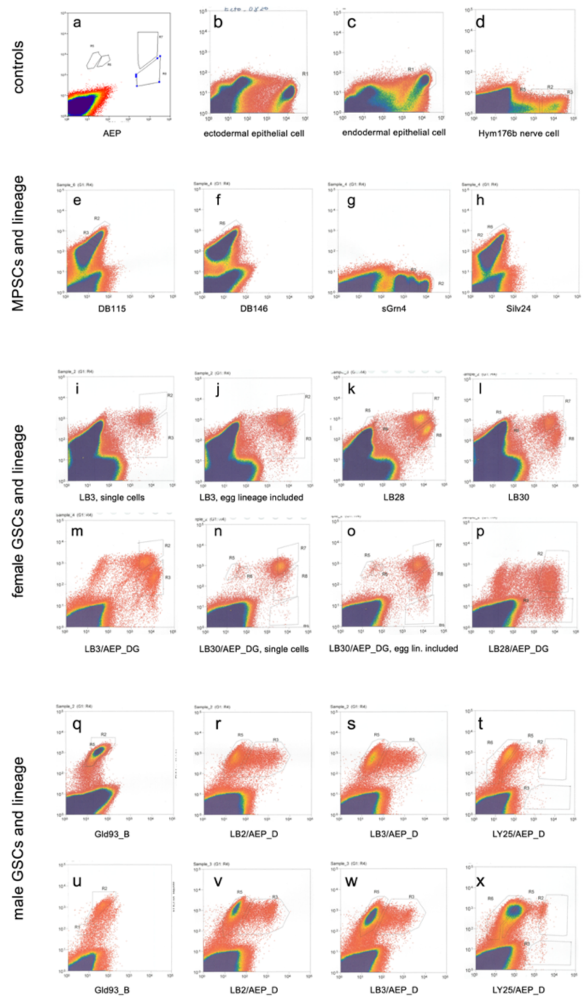


**Fig. S5.** Isolation of *Hydra* cell types by FACS. Sorting for sorted cells by using the following strains (see also Table S1): (a) The original non-transgenic strain *Hydra vulgaris* AEP; (b) ectodermal epithelial cells, strain A16 (*actin::eGFP*); (c) endodermal epithelial cells, strain A17 (*actin::eGFP*); (d) nerve cells, strain Toshi_B (*Hym176B::eGFP*); (e MPSCs and lineage, strain DB115 (*Cnnos1::DsRED; DM5::eGFP*); (f) MPSCs, strain DB146 (*Cnnos1::DsRED; DM5::eGFP*); (g) MPSCs and lineage, strain sGrn4 (*Cnnos1::eGFP*); (h) MPSCs and lineage, strain Silv.24 (*Cnnos1::DsRED*; (i) ♀GSCs (single cells), strain LB3 (*Cnnos1::DsRED; HybolF1::eGFP*); (j) ♀GSCs and lineage, strain LB3 (*Cnnos1::DsRED; HybolF1::eGFP*); (k) ♀GSCs and lineage, strain LB28 (*Cnnos1::DsRED; HybolF1::eGFP*); (l) ♀GSCs and lineage, strain LB30 (*Cnnos1::DsRED; HybolF1::eGFP*); (m) ♀GSCs and lineage, strain LB3/AEP_DG (*Cnnos1::DsRED; HybolF1::eGFP*); (n) ♀GSCs single cells, strain LB3/AEP_DG single cells (*Cnnos1::DsRED; HybolF1::eGFP*); (o) ♀GSCs, strain LB3/AEP_DG  single cells & lineage (*Cnnos1::DsRED; HybolF1::eGFP*); (p) ♀GSCs, strain LB28/AEP (*Cnnos1::DsRED; HybolF1::eGFP*); (q) ♂GSCs, strain Gold93B (*Cnnos1::DsRED*; *DM5::eGFP*); (r) ♂GSCs & lineage, strain LB2/AEP_D chimera (*Cnnos1::DsRED; HybolF1::eGFP*); (s) ♂GSCs & lineage, strain LB3/AEP_D chimera (*Cnnos1::DsRED; HybolF1::eGFP);* (t) ♂GSCs & lineage strain LY25/AEP_DG chimera *(Cnnos1::DsRED; DM5::eGFP; HybolF1::eGFP);* (u) ♂GSCs, strain Gold93B (*Cnnos1::DsRED*; *DM5::eGFP*); (v) ♂GSCs & lineage, strain LB2/AEP_D chimera (*Cnnos1::DsRED; HybolF1::eGFP*); (w) ♂GSCs & lineage, strain LB3/AEP_D chimera (*Cnnos1::DsRED; HybolF1::eGFP);* (x) ♂GSCs & lineage strain LY25/AEP_DG *(Cnnos1::DsRED; DM5::eGFP; HybolF1::eGFP)*.

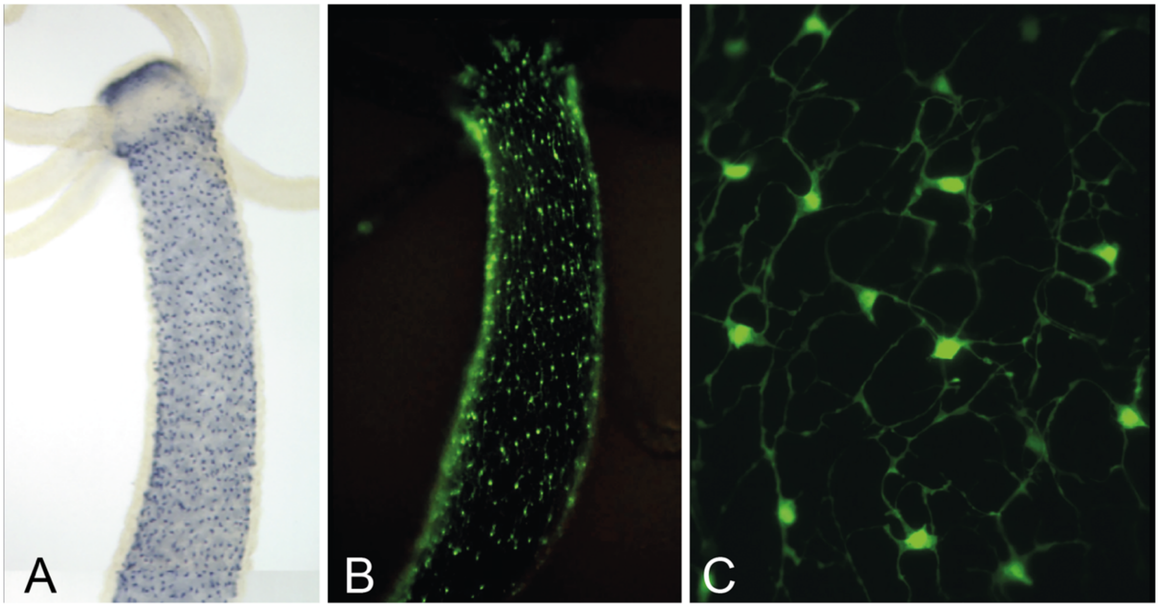


**Fig. S6.** Polyp transgenic for Hym176B neurons. (a) ISH shows the expression of Hym176B transcript along the upper body column of a polyp. (b) Distribution of Hym176B::GFP along the upper body column of a polyp and (c) network of Hym176B::GFP at higher magnification

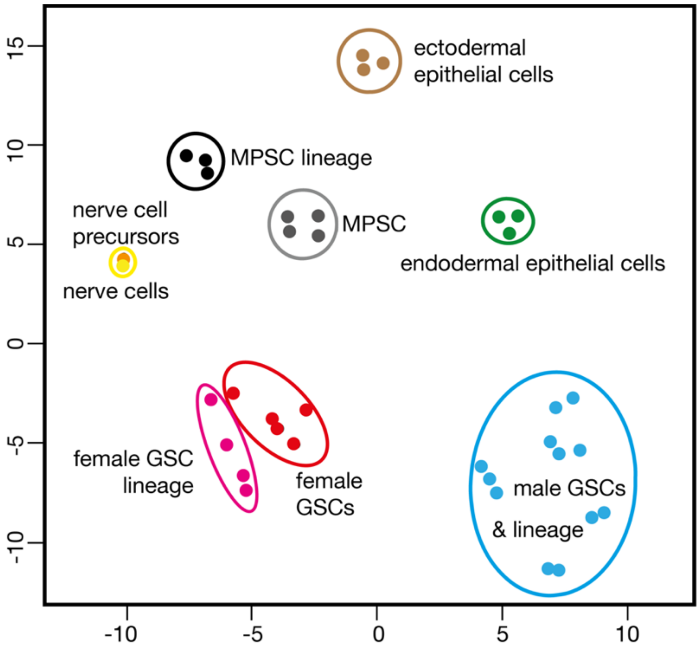


**Fig. S7.** t-SNE assay for homogeneity of cell type categories. Normalized and rlog transformed data were clustered and visualized by a modified t-SNE (T-distributed Stochastic Neighbor Embedding) algorithm using the R implementation (Rtsne).

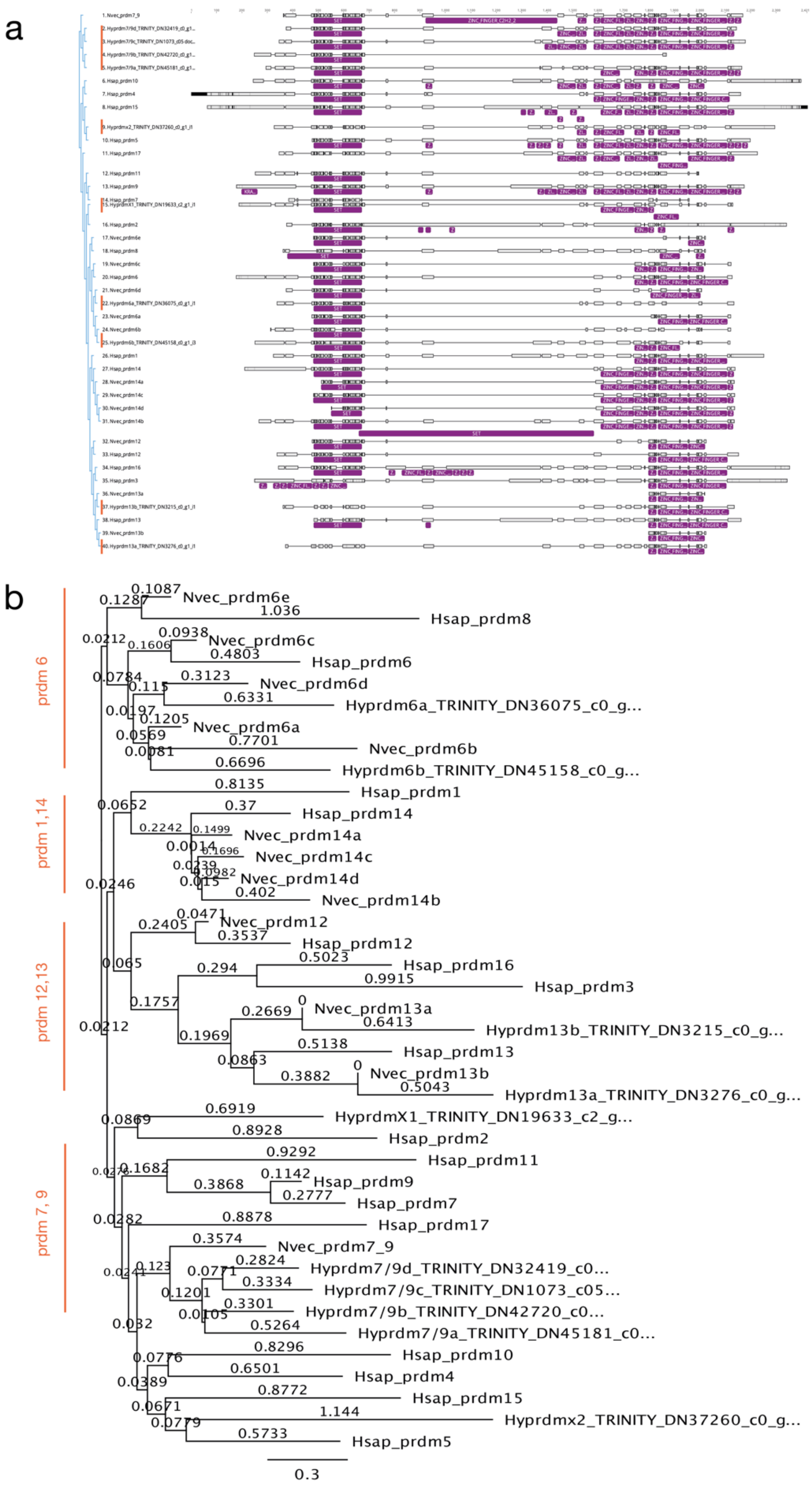


**Fig. S8.** Molecular phylogeny of Hydra PR domain zinc finger proteins (prdm). (a) Multiple alignment with Human, Hydra and Nematostella sequences; (b) NJ-tree of ortholgous prdms.

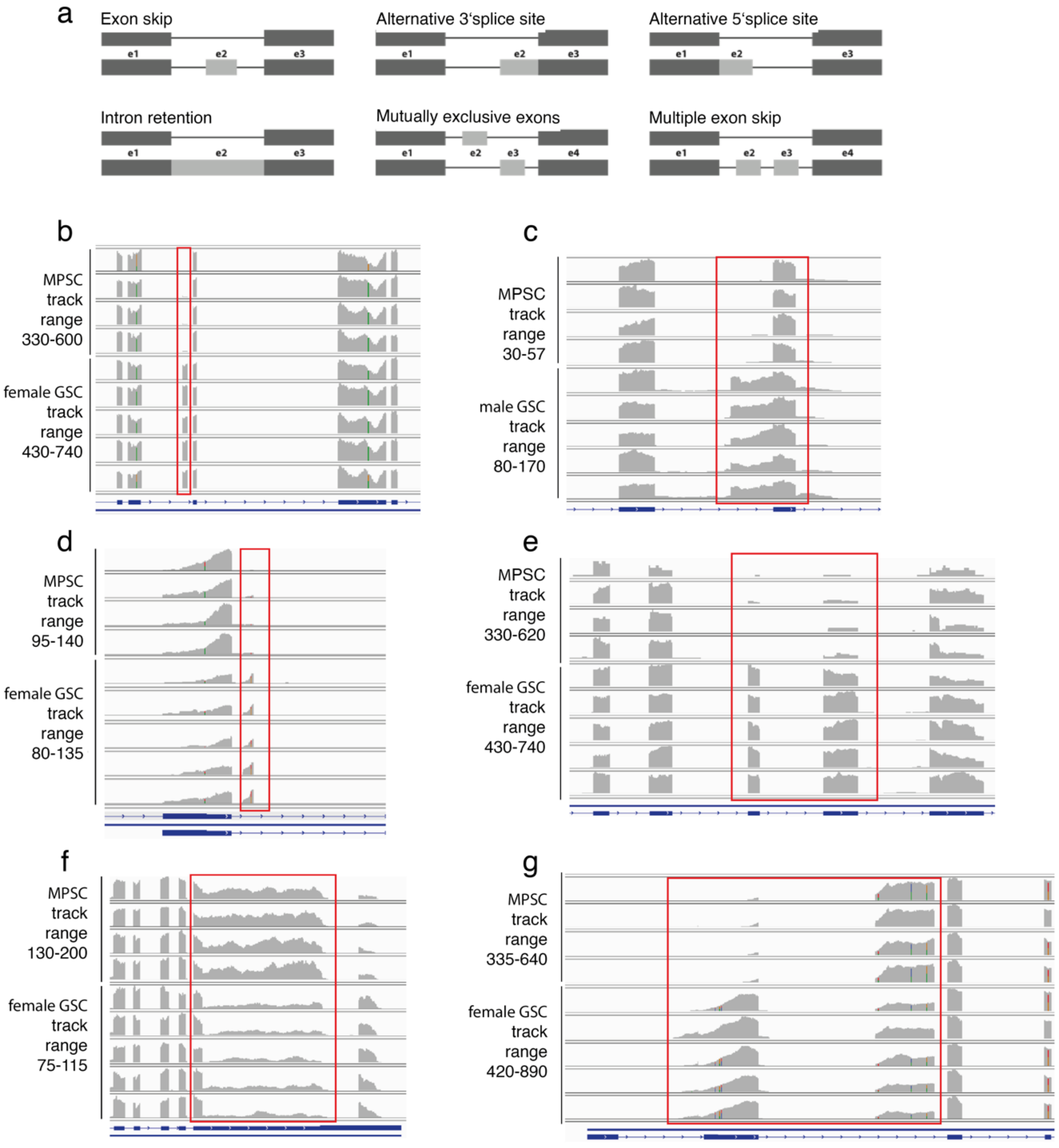


**Fig. S9**. Spladder identified alternative splicing events and example genes, adjusted p-value < 0.0001. (a) Spladder alternative splicing event types. (b-g) Example genes on IGV. (b) Exon skip, MPSC/female GSC, *BAZ2B* (HVAEP11.G020765); (c) Alternative 3' splice site, MPSC/male GSC, *KIF1A* (HVAEP15.G027788); (d) Alternative 5' splice site, MPSC/female GSC, *HMCN1* (HVAEP11.G020663); (e) MPSC/female GSC multiple exon skip; *ANR31* (HVAEP15.G027795); (f) Intron retention, MPSC/female GSC, *DCLK2* (HVAEP12.G021315); (g) MPSC/male GSC mutually exclusive exons, *HNRPL* (HVAEP10.G019194).

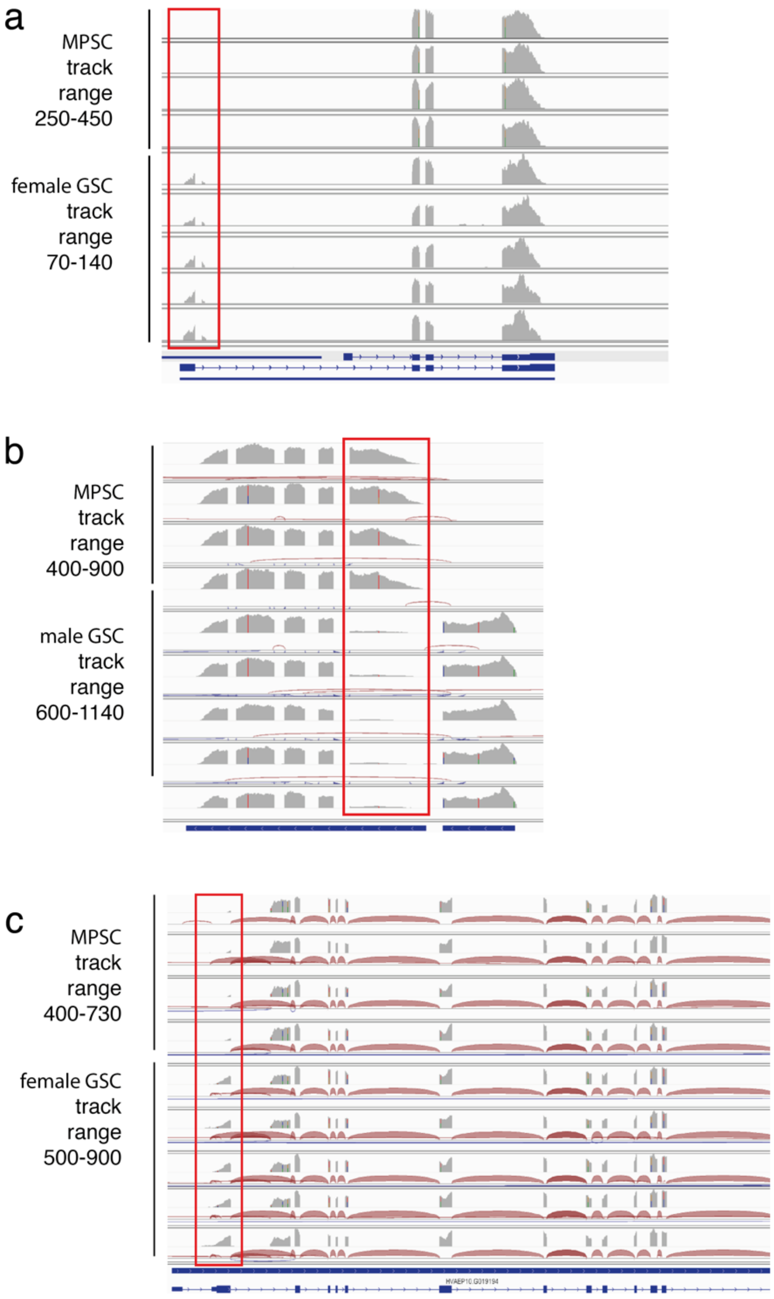


**Fig. S10.** Splicing patterns of key cell type regulators**.** Splicing patterns of Otx2, Snf5 and Hnrnpl revealed splicing between MPSC/female GSC and MPSC/male GSC. (a) Key germline regulator *Otx2* (HVAEP6.G011272, TRINITY_DN38831) revealed alternative splicing pattern between MPSC and female GSC. (b) Metazoan cell type differentiation regulator *Snf5* (HVAEP10.G018475, TRINITY_DN34020) reveals alternative splicing isoform in male GSC. (c) Epigenetic factor Hnrnpl (HVAEP10.G019194, TRINITY_DN46875) revealed alternative splicing pattern between MPSC and female GSC

### TABLES

**Table S1**. Strains used in this study.

**Table1: Strains used in this analysis**

| Strains | Reporter | Strain Description | Transgene(s) | GSC regeneration experiment | Sorted cell type or tissue: ectodermal epithelial cells | endodermal epithelial cells | MPSCs | MPSC lineage | ♀ GSCs | ♀ GSC lineage | ♂ GSCs | ♂ GSCs | ♂ GSC lineage | nerve cells (eGFP-strong) | nerve cells (eGFP-weak) | whole tissue* as, sm, sf, sh |
|---|---|---|---|---|---|---|---|---|---|---|---|---|---|---|---|---|
| A14 | G | Founder (Chimera) | (actin::eGFP) / ectodermal epithelial cells | | • | | | | | | | | | | | |
| A16 | G | Founder (Chimera) | (actin::eGFP) / endodermal epithelial cells | | | • | | | | | | | | | | |
| DB115 | D** | Founder (Born Chimera: Tg only in MPSCs) | (Cnnos1::DsRED; DM5::eGFP) / MPSC | | | | • | • | | | | | | | | |
| DB146 | D** | Founder (Born Chimera: Tg only in MPSCs) | (Cnnos1::DsRED; DM5::eGFP) / MPSC | | | | • | | | | | | | | | |
| sGrn4 | G | Founder (Born Chimera: Tg only in MPSCs) | (Cnnos1::eGFP) / MPSC | ▲ | | | • | • | | | | | | | | |
| sGrn4$^{regenerate}$ | | Founder (GSCs are regenerated from TgMPSCs) | | △ | | | | | | | | | | | | |
| Silv.24 | D | Founder (Born Chimera: Tg only in MPSCs) | (Cnnos1::DsRED) / MPSC | ▲ | | | • | • | | | | | | | | |
| Silv24$^{regenerate}$ | D | Founder (GSCs are regenerated from TgMPSCs) | (Cnnos1::DsRED) / (MPSC & GSCs) | △ | | | | | | | | | | | | |
| Silv24$^{regenerate}$-6 | D | F1 (Silv24$^{regenerate}$ x AEP) | (Cnnos1::DsRED) / (MPSC & GSCs) | | | | | | | | | | | | | |
| LWA | G | Founder | HybolF1::eGFP/ EgSC | | | | | | | | | | | | | |
| LWA-4 | G | F1 (LWA x AEP) | HybolF1::eGFP/ EgSC | | | | | | | | | | | | | |
| LWA-16 | G | F1 (LWA x AEP) | HybolF1::eGFP/ EgSC | | | | | | | | | | | | | |
| LB2 | D; G | F2 (Silv24-6 xLWA-4) | Cnnos1::DsRED; HybolF1::eGFP | | | | | | | | | | | | | |
| LB3 | D; G | F2 (Silv24-6 xLWA-16) | Cnnos1::DsRED; HybolF1::eGFP | | | | | | • | • | | | | | | |
| LB28 | D; G | F2 (Silv24-6 xLWA-16) | Cnnos1::DsRED; HybolF1::eGFP | ▲ | | | | | • | • | | | | | | |
| LB30 | D; G | F2 (Silv24-6 xLWA-4) | Cnnos1::DsRED; HybolF1::eGFP | | | | | | • | • | | | | | | |
| Gold93B | D*** | Founder (Born Chimera: Tg only in SpSCs) | (Cnnos1::DsRED; DM5::eGFP) / SpSC | | | | | | | | | • | | | | |
| LY25 | D; G | F1 (Gold93B x LWA-16) | Cnnos1::DsRED; DM5::eGFP; HybolF1::eGFP | | | | | | | | | | | | | |
| LB3 / AEP_DG | D; G | F2-Chimera, Female | (Cnnos1::DsRED; HybolF1::eGFP) / EgSC | | | | | | • | | | | | | | |
| LY25 / AEP_DG | D; G | F2-Chimera, Female | (Cnnos1::DsRED; DM5::eGFP; HybolF1::eGFP) / EgSC | | | | | | • | • | | | | | | |
| LB2 / AEP_D | D**** | F2-Chimera, Male | (Cnnos1::DsRED; HybolF1::eGFP) / SpSC | | | | | | | | ○ / • | ○ / • | ○ / • | | | |
| LB3 / AEP_D | D**** | F2-Chimera, Male | (Cnnos1::DsRED; HybolF1::eGFP) / SpSC | | | | | | | | • | • | • | | | |
| LY25 / AEP_D | D | F1-Chimera, Male | (Cnnos1::DsRED; DM5::eGFP; HybolF1::eGFP) / SpSC | | | | | | | | | • | • | | | |
| Toshi_B | G | F1 | Hym176B::eGFP | | | | | | | | | | | • | • | |
| AEP | - | Original Strain | - | | | | | | | | | | | | | • |

In all cases, heads and feet were removed before tissue disociation.
For sexual tissue, only sexually differentiating parts of the bodies were used.
▲: strains used for GSC-regeneration
△: strains resulted from GSC-regeneration experiment
●: Cells were collected from animals cultured at 18˚C
○: Cells were collected from animals cultured at 15˚C to increase the stem cell fraction.
* In addition we sequenced tissue from asexual (as) polyps, sexual gonochoristic polyps undergoing oogenesis (sf) or spermatogenesis (sm), and hermaphroditic polyps undergoing oogenesis and spermatogenesis (sh).
** Only MPSCs were transgenic in this strain. Therefore the male-specific DM5::eGFP is not detectable in any stage of sexual differentiation (DM5/Dmrt1 is the so far undescribed hydra ortholog of transcription factors that act in bilaterian gonad development (37, 38).
*** SpSCs in this strain is double transgenic, but DM5::eGFP was undetecttable even with FACS.
**** leaky HybolF1::eGFP

**Legends to Table S2-12**.

**Table S2.** Cell type specificity of all analysed *Hydra* genes (submitted to GEO/GSE163910). Several specific tables were extracted from this comprehensive table:  Supplementary Tables 3-5 (male and female GSCs), Supplementary Tables 6-9 (molecular factors), Supplementary Table 10 (MPSCs), Supplementary Table 11 (alternative splicing in in MPSCs and in male and female GSCs), Supplementary Table 12 (functional annotation of gene in MPSCs and GSCs alternatively spliced)

**Table S3**. Genes enriched in male and female GSCs.

**Table S4**. Genes enriched in female GSCs.

**Table S5**. Genes enriched in male GSCs.

**Table S6**. Genes enriched in MPSCs.

**Table S7**. Cell type distribution of epigenetic factors.

**Table S8**. Cell type distribution of Bmp signalling.

**Table S9.** Cell type distribution of Wnt signalling.

**Table S10**. Cell type distribution of transcription factors.

**Table S11**. Alternative splicing in MPSCs and GSCs**.**

**Table S12**. Annotation of alternatively spliced genes.
